# Correlating Structure and Rheology in Ionically Crosslinked Alginate Biopolymer Hydrogels - A Case for Why “Less” can be “More”

**DOI:** 10.64898/2026.07.23.740415

**Authors:** Vinay Kopnar, Peter S Sherin, Sarah Graham, Harriet Fyfe, Reyna Garcia-Gonzalez, Adam O’Connell, Natasha Shirshova, Anna Barnard, John Girkin, Marina K Kuimova, John Bothwell, Anders Aufderhorst-Roberts

**Author notes:** AA-R, MKK, JG, and JB conceptualized the study. VK, PS, SG, HF and RGG carried out the experimental work and analysed the data. VK, PS and AA-R prepared the original draft. NS, AO, AB, MKK, JG, JB and AA-R supervized the work. PS, SG, AA-R, MKK, JG and JB reviewed and edited the article. There are no competing interests.

## Abstract

We probe the structural and rheological properties of ionically crosslinked alginate, a model biopolymer hydrogel, using microscopy, rheology and viscosity dependent molecular probes. This combination of techniques enables the quantification and correlation of microstructure, microviscosity, bulk viscoelasticity and yielding dynamics. By adjusting the stoichiometric ratio, R, between the alginate biopolymer and its cation crosslinks, we observe a transition from a homogeneous network-like structure to a coarse bundle-like structure at high (R*>*0.67) stoichiometric ratio. Intriguingly, these bundle-like structures have distinct and counter-intuitive rheological properties. Using molecular probes, we observe a continuous decrease in microviscosity that is correlated with a decrease in bulk elastic modulus and an increase in energy dissipation. This is accompanied by a transition in the hydrogel yielding under strain from a sharp, well-defined yield point to a continuous ductile-like yielding. We ascribe these surprising transitions to the looser intermolecular interaction between alginate biopolymers in the bundle-like state, as previously predicted by x-ray scattering experiments. These findings reveal new and counter-intuitive structure-property relations that demonstrate high crosslink concentration does not necessarily translate to optimal mechanical performance.

**Significance:** Alginate is a polysaccharide biopolymer naturally found in brown seaweed cell walls. Extracted alginate forms ionically crosslinked hydrogels that are strong, flexible and increasingly valuable in the biomedical, packaging and food industries. A large part of the utility of these hydrogels stems from the ease with which their mechanics can be tuned through adjusting the stoichiometry between alginate and its ionic crosslinks. However, little is known about how the material properties of alginate hydrogels, in particular their rheology and dynamics, are affected by microscale structural transitions at high stoichiometric ratios. Here, we use a multi-modal approach to describe and correlate hydrogel material properties and to demonstrate that increased polymer crosslinking can, counterintuitively, sometimes weaken hydrogel performance.

---

Charged polymeric hydrogels derived from natural polysaccharides offer exciting platforms for soft structured materials because of their biodegradability, abundance, and novel rheological properties. The polysaccharides found in the cell walls of the macroalgae, or seaweeds, are of particular commercial interest as they can be readily crosslinked by cations, leading to highly tunable mechanical properties (1). These crosslinked networks impart mechanical stability to cell walls in nature and also have promising futures in food (2), packaging (3), and biomedical applications (4, 5). In the current study, we focus on alginate, an extensively-studied polysaccharide that is extracted from brown seaweed and that undergoes gelation in the presence of, most commonly, calcium ions. X-ray scattering studies on alginate (6) have shown that varying the ratio of calcium ions to polymer can drive the formation of a range of molecular associations and hydrogel microstructures. However, the relationship between these microstructures and the bulk rheological properties of their hydrogels remains an open question that bridges multiple length scales. Answering this question is of central importance in understanding both the native mechanical properties of marine cell wall biopolymers and the formulation of manufactured materials from sustainable feedstocks.

Chemically, alginate is a heterogeneous block co-polymer: it comprises a linear chain of mannuronic acid (M units) and guluronic acid (G units), but the repeat pattern and length of these M and G units varies between brown algal species (7). Mechanistically, alginate gelation is mediated by divalent ions that preferentially join repeating G units on separate chains into a dimeric conformation (Figure 1(a)), leading to the formation of a so-called ‘egg-box’ dimer. The structure of an alginate hydrogel depends on the stoichiometric ratio, R, which describes the ratio of divalent cations to G units.

**Fig. 1.**
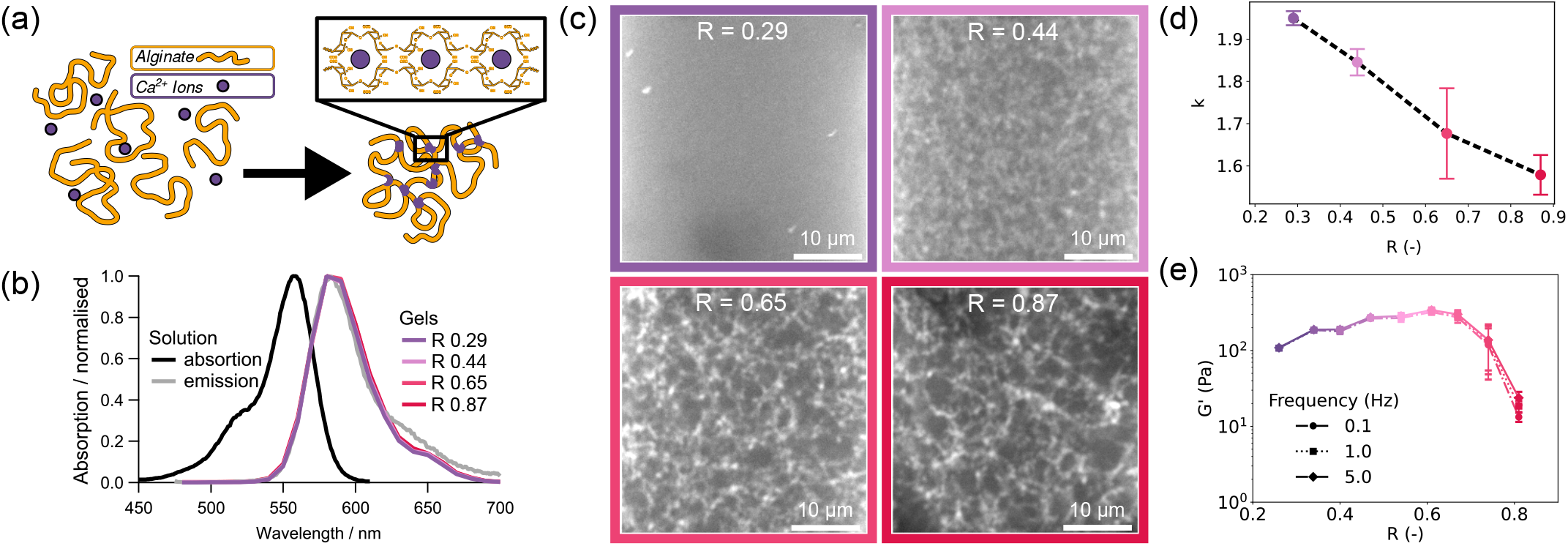
Confocal fluorescence micrographs of alginate structures at variable stoichiometric ratio R. (a) Alginate hydrogels form when divalent ions bind to the guluronic acid groups on the biopolymer backbone, leading to a crosslinked structure known as an egg-box dimer; (b) Absorption spectrum of AL510 in aqueous solution (in black) (0.15 µg/ml) and emission spectra (in gray) in aqueous solution (0.15 µg/ml) and in hydrogels (13.5 mg/ml); (c) Fluorescence intensity images obtained with AL510 hydrogels recording the emission in the 540-680 nm range following the two-photon excitation at 930 nm; (d) Analysis of microstructure within AL510 hydrogels, presented in (c) using a fractal model; (e) shows the evolution of storage modulus (G’) under small-amplitude oscillatory shear at three frequencies of oscillation with increasing *R*.

At the lowest stoichiometric ratios (*R* < 0.25), precursor structures are believed to form that comprise either mono-complexes of cations bound to a single chain (8) or so-called ‘tilted’ dimers that link two separate chains. (9). For higher ratios of 0.25 < *R* < 0.55, more rigid egg-box dimer structures form (6), leading to unambiguous gelation. At still higher concentrations (*R >* 0.55), the consensus is less clear. Earlier studies suggested that egg-box dimers form higher order structures via lateral associations (6), driven by the neutralization of the carboxyl groups on the M units of alginate chains (10, 11). The resulting structures have been observed in solution and are thought to be crosslinked through noncovalent interactions such as hydrogen bonding and nonspecific binding of ions to M units (12, 13). Here, we describe these structures as “bundle-like” (14) to distinguish them from the network-like structures commonly assumed to support hydrogel formation but other names, including egg-box multimers (6, 15) and egg-box junctions, have also been used (16). While the formation of these bundle-like structures is not disputed, it remains entirely unclear how these structures influence the alginate’s material properties. There are some hints that the alginate hydrogel elastic modulus scales differently with R, in the low and intermediate Ca^2+^ regimes (17), that the solutions of bundle-like alginates have lower viscosities (13) and that distinct relaxation timescales begin to emerge at intermediate R values (16). However, a direct relation between structure and rheology has yet to be identified and the biophysically crucial question of how these structural changes affect the hydrogel’s microstructural breakdown under shear remains unanswered.

Accordingly, here we address those questions by adopting a multi-length-scale approach. First, we directly visualize the crosslinking of alginate using fluorescently labelled alginate, synthesizing the hydrogels at selected *R* values that encompass the full spectrum of alginate microstructure regimes reported in the literature. Next, we apply fluorescent lifetime techniques to quantify the apparent molecular viscosity in the hydrogel network at different stoichiometric ratios. Hydrogel rheology under these conditions is then probed using standard small amplitude oscillatory shear. Finally, the microstructural breakdown of these different alginate hydrogels is probed using large amplitude oscillatory shear (LAOS) rheology, a data-rich rheological technique that entails applying sinusoidal shear strain cycles of increasing amplitude. By applying this multi-modal approach to alginate hydrogels with different microstructures, we establish a structure-property relationship for the alginate hydrogels that spans length scales. With the growing use of alginate in industrial applications as a sustainable alternative to petrochemical plastics, our results have implications for formulation challenges during product development, where the relationship between crosslinking and material properties plays a crucial role. The wide seasonal and species-specific range of alginate compositions seen in living cell walls also suggests physiological roles for these structure-property relationships in protecting algal cells from mechanical stress (18).

## Results and Discussion

### Alginate hydrogels develop bundle-like structures at high R

Alginate hydrogels were prepared using the so-called internal gelation approach,(19) in which calcium ions are released internally in an alginate solution through gradual hydrolysis. The resulting release of calcium ions leads to the gradual formation of egg-box dimers over time as shown in figure 1(a). Our key design constraint is the stoichiometric ratio, R,(6, 10) which describes the ratio of divalent calcium ions to G units and acts as a predictor of the resulting stiffness and failure properties of hydrogels (20).

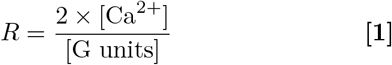

We selected crosslinking stoichiometric ratios in the range 0.3 *<*R*<* 1.0. The lower bound of this range corresponds to the expected critical crosslinking threshold of alginate (R = 0.25) under dilute conditions (6) and the upper bound corresponds to the conditions under which bundle-like structures have previously been experimentally observed (R > 0.55). (21).

We probe the microstructure of the resulting hydrogels using an rhodamine B (RhB)-labelled sodium alginate (AL510) where RhB isothiocyanate is joined to the sodium alginate backbone via an ethylenediamine spacer (SI Appendix,Fig S1). We first verify that RhB does not interfere with the crosslinking of alginate chains. The hydrogels were prepared with a range of stoichiometric ratios and exhibited similar transparency and color (SI Appendix, Fig. S1). Furthermore, the shape of the RhB emission band (Figure 1(b)) remained unchanged with increasing *R*. These observations indicate that RhB does not undergo any chemical modification and is not involved in the crosslinking of polysaccharide chains. The evolution of observed microscale patterns of hydrogels as viewed under confocal microscope with variations in *R* is given in Figure 1(c). At low *R* ∼ 0.29, the microstructure appears diffuse, indicating structures that are below the diffraction limit. At increasing R, increasingly coarse structures are observed such that at the highest *R* = 0.87, the hydrogel mesh appears open with the clear presence of thick bundle-like structures. To quantify these observed patterns, we calculated their fractality using a box-counting algorithm (22) (SI Appendix, Fig. S3 & S4). The output of this algorithm is the two-dimensional network fractality factor k, which has a theoretical range of 0 *< k <* 2, the limits of which correspond to a homogeneously filled environment (k = 2) and discrete point-like structures (k = 0) respectively. Applying this algorithm reveals an unambiguous decrease in k with *R* (Figure 1(d) and SI Appendix, Fig. S5), indicating that the fractality of the microstructures increases with the crosslink stoichiometric ratio. The increase in fractality demonstrates that the bundle-like structures previously observed under dilute conditions (23) also manifest within alginate hydrogels. To investigate the rheological properties of these alginate hydrogels, we use unlabeled alginate powder to synthesize different hydrogels varying *R* from 0.27 to 0.81, maintaining the alginate concentration and thus the concentration of G units. We perform small amplitude oscillatory shear imparting sinusoidal deformation with low amplitude (*γ*_0_ = 0.01) that is well within the LVE regimes of the hydrogels. Figure 1(e) presents the storage modulus (G’), which exhibits a weak dependence on *R* within the range 0.27 *<*R*<* 0.47, indicating an increase in crosslink density formed between G units and Ca^2+^. For *R >* 0.47, the modulus remains approximately constant until *R* reaches 0.67, highlighting that all the possible egg-box dimers have been formed. Within this range we may assume that G’ is inversely related to the average chain length between the crosslinks, as predicted by rubber elasticity theory (24) The upper limit of this regime is consistent with alginate stoichiometry since *R* ∼ 0.47 - 0.54 corresponds to the approximate limit of stoichiometric balance in which one Ca^2+^ ion binds to four G units (Figure 1(a)). On the microstructural level, the limit *R >* 0.55 also corresponds to the approximate point at which bundle-like structures are expected to form (6). For *R >* 0.67, the value of G’ becomes inversely dependent on R. Drawing again on rubber elasticity theory, this indicates some form of systematic change in structure for *R* ≥ 0.67, that would necessitate an increase in the distance between the crosslinks in the network. This is remarkably consistent with the microstructural data shown in figure 1 (c) and fractality analysis in figure 1 (d) which show a more porous network with bundle-like structures.

### Bundle-like hydrogels exhibit lower viscosities at both microscopic and macroscopic length scales

Having demonstrated the presence of bundle-like structures and quantified the resulting changes in rheology on the macroscopic scale, we now examine the rheological properties on the local microscopic scale. We construct hydrogels from RhB-labeled alginate (AL510) and carry out spatially resolved FLIM measurements. The RhB fluorophore is known for its sensitivity to the local microviscosity(25, 26) which affords the opportunity to provide additional information about the hydrogel micro-organization. Small fluorescent probes sensitive to viscosity, so-called molecular rotors, are commonly used to determine the viscosity at the microscale by monitoring their response in both fluorescence intensity and lifetime (27). The fluorescence lifetime of RhB is independent of its concentration and RhB does not interact with alginate (25) making it ideal to sense the variations in local viscosity induced by alginate crosslinking.

Figure 2(a) shows the maps of fluorescence lifetime distribution of alginate hydrogels prepared using four selected values of R, showing a consecutive decrease of RhB lifetime with the increase of *R* value. Further evidence for this decrease is given by recording decay traces showing steepening slope as R increases from 0.29 to 0.87 in Figure 2(b) and histograms of lifetime distributions in Figure 2(c). Notably, there is no correlation between intensity and lifetime data (SI Appendix, Fig. S6). This indicates that the spatial variation in dye concentration (fluorescence intensity) is independent of the local micro-viscosity (fluorescence lifetime). Figure 2(d) presents the lifetime of two AL510 solutions. As the concentration is increased fourfold from 54 mg/ml to 0.15 *µ*g/ml, the lifetime is observed to decrease by only 2%, demonstrating conclusively that the lifetime is independent of AL510 concentration.

**Fig. 2.**
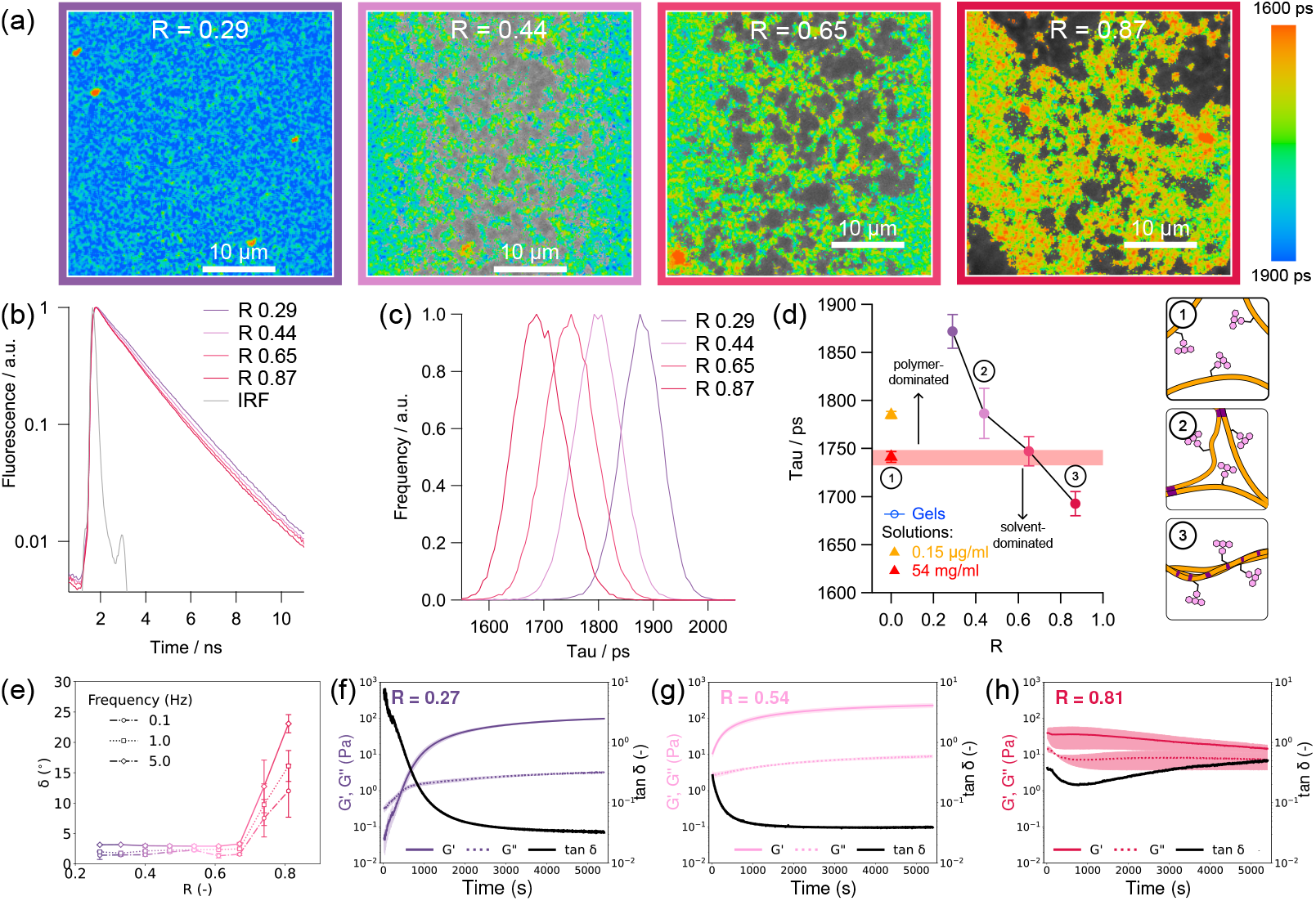
Fluorescence lifetime characterization of RhB used to label alginate chains of AL510 hydrogels and hydrogel formation kinetics of alginate hydrogels of different *R* in the corresponding measurement range. (a) FLIM images obtained with AL510 hydrogels recording the emission in the 540-680 nm range following the two-photon excitation at 930 nm; The color bar on the right shows the fluorescence lifetime. (b) Decay traces averaged over the frame. The IRF curve represents the Instrument Response Function which was deconvolved to give accurate lifetimes; (c) Lifetime distributions for FLIM data in (a); (d) *R*-dependence for the averaged lifetimes from (c) and AL510 in aqueous solution at low and high concentrations (0.15 µg/ml and 54 mg/ml, orange and red triangles, respectively) presented as *R*= 0. Square boxes show representation of the interaction and lifetime of RhB with alginate in the presence and absence of Ca^2+^ ions; (e) shows the evolution of phase angle, *δ*, under small-amplitude oscillatory shear at three frequencies of oscillation with increasing *R*; (f), (g), and (h) present the evolution of G’, G” and tan *δ* with time during the gelation of alginate at *R* = 0.27, 0.54, and 0.81 respectively.

As shown in figure 2(d) the corresponding fluorescence lifetime of alginate hydrogels decreases systematically as *R* increases. Since the alginate concentration does not appear to affect the RhB lifetime, the lifetime changes could not be related to alginate aggregation-induced processes and can only be related to interaction of RhB with its viscous environment. As the RhB is constrained within a crosslinked alginate network, we intuitively expect the apparent viscosity of the hydrogel to be higher than that of the alginate solutions, corresponding to a higher fluorescence lifetime. Fig 2(d) shows that this is indeed the case within the limit *R* < 0.65. However For *R* ≥ 0.65, corresponding to the conditions in which bundle-like structures form, the RhB lifetime in the hydrogel is lower than the equivalent lifetime in solution. Counterintuitively, this indicates that the microviscosity within the hydrogel network is lower than in solution. We rationalize this by considering that the formation of bundle-like structures would expose RhB to the aqueous environment (③ in Figure 2(d)). To test this assumption, we recorded the viscosity calibration curves for RhB as a free dye and compared this to measurements of RhB-labeled alginate in a set of binary mixtures of water and glycerol (detail in SI Appendix). Both samples demonstrate viscosity dependence, which is somewhat reduced for AL510, possibly due to steric hindrance induced by polysaccharide chains (Figures 2(d) and SI Appendix, Fig. S7). As a final test we also considered that the water-induced quenching of fluorescence lifetime, which is a well-known phenomenon for many organic dyes, could play a role in the observed changes in fluorescence lifetimes (28). To verify the presence of any solvent quenching effect for RhB, we carried out a control experiment in a set of alcohols (SI Appendix, Fig. S9 and S10). Our results (SI Appendix, Fig. S10 and Table S1) showed a consecutive decrease in RhB lifetime from tert-butanol to water, which correlates with both the solvent’s ability to donate hydrogen bonds (29) and the density of OH bonds in solvents. Considered together, the observed decrease in lifetimes for AL510 in hydrogels can be interpreted as (1) a reduction in RhB microviscosity combined with (2) a partial quenching of RhB emission by water molecules. Both phenomena are consistent with higher solution exposure of RhB due to the progressive formation of bundle-like structures with increasing *R* (Figure 2(d)).

Having probed the microviscosity of the hydrogels, we now quantify their macroscopic viscosity by using oscillatory shear rheology to probe the phase angel *δ* = tan ^-1^ (G”/G’), which is a measure of a material’s propensity to dissipate energy. Similar to Figure 2(d), *δ* depicts two regimes. For *R* ≤ 0.67, *δ* remains close to zero, suggesting that the energy is predominantly stored elastically. For *R >* 0.67, *δ* is observed to increase sharply, indicating a disproportionate increase in energy dissipation. This is in close agreement with the decrease in the lifetimes of RhB observed at the corresponding range of *R*. Taken together, this demonstrates that the emergence of bundle-like structures for *R >* 0.67 leads to hydrogels that store and dissipate energy in way that is entirely distinct from those formed when *R* ≤ 0.67.

### Bundle-like hydrogels exhibit non-monotonic gelation kinetics

To further understand the various structures formed in alginate hydrogels we also probe their kinetics of gelation. Fig. 2(f)-(h) shows three exemplar plots of the evolution of G’ and G” as the hydrogels are formed in-situ. For all hydrogels, a rapid increase in G’ and G” is observed from the earliest times of gelation, indicating the rapid crosslinking of alginate chains. At the lowest *R* values, this increase is sufficiently gradual that we observe a crossover between G’ and G”, which is generally considered a crude estimate of the critical gel point (30). The crossover point depends on the binding coefficient of calcium to alginate (19) such that the associated critical timescale, known as the Smoluchowski doubling time, is inversely proportional to *R* (31). The post-gelation evolution of G’ and G” for hydrogels with the highest *R* (≥ 0.67) is distinctly different. After gelation, there is a decrease in both moduli suggesting some form of secondary structural evolution post-gelation. Further evidence of this can be seen by plotting the value of tan *δ* = G”/G’, which is a measure of a material’s ability to dissipate energy. For *R* < 0.67, tan *δ* decreases steadily, suggesting a continuous evolution structure through increments in the elasticity of hydrogels due to coarsening of the egg-box dimers. By contrast at high *R* (≥ 0.67), the value of tan *δ* first decreases and then increases. The initial decrease in tan *δ* is consistent with rapid gelation, while the subsequent decrease is often observed in hydrogels involving physical crosslinks, where it is generally thought to relate to post-gel structural changes in the aggregates or bundles formed during the gelation process (32–34).

### Bundle-like hydrogels exhibit ductile yielding

So far we have focused on the linear viscoelastic properties of alginate hydrogels. To understand how these hydrogels undergo structural breakdown, we now turn to nonlinear rheology by applying large oscillatory strains beyond the linear viscoelastic regime (35–37). We use LAOS to apply sinusoidal oscillatory cycles of increasing strain amplitude (38). These responses elucidate fundamental material behaviors such as strain-stiffening and yielding mechanisms, which offer a window into alginate microstructure. We begin by examining the average changes in G’ and G” of the hydrogels with increasing *γ*_0_. Fig. 3 shows this progression for hydrogels with different *R*. Neither modulus shows any change at low *γ*_0_, highlighting the LVE regime of the hydrogels where all the deformation is recovered by the hydrogels once any applied strain is removed. For hydrogels with *R* < 0.67, the LVE regime extends until *γ*_0_ ∼ 0.4 (Fig. 3(a),(c), &(e)) while for hydrogels with *R* ≥ 0.67, the LVE regime becomes narrower (Fig. 3(h) & (j)). A marked difference is observed in the progression of both the moduli when hydrogels enter the non-LVE regime. The value of G” exhibits a sharp rise and more gradual fall for hydrogels with *R* < 0.67. This overshoot in G” reports the unrecoverable deformation with increasing *γ*_0_ (39) and is driven by a breakdown of microstructural constraints(40, 41). This overshoot reduces to the point of being negligible with increasing R. G’, on the other hand, declines for all the hydrogels in the non-LVE regime irrespective of *R*. However, there is a sharp difference in the nature of the decline. For hydrogels with *R* < 0.67, the decline in G’ is sharp as opposed to the decline in G’ for hydrogels with *R* ≥ 0.67, when the slope of decline is gradual. These differences in G’ and G” suggest that the hydrogels with *R* < 0.67 undergo a sudden brittle yielding transition while hydrogels with higher *R* undergo a gradual ductile yielding transition.

**Fig. 3.**
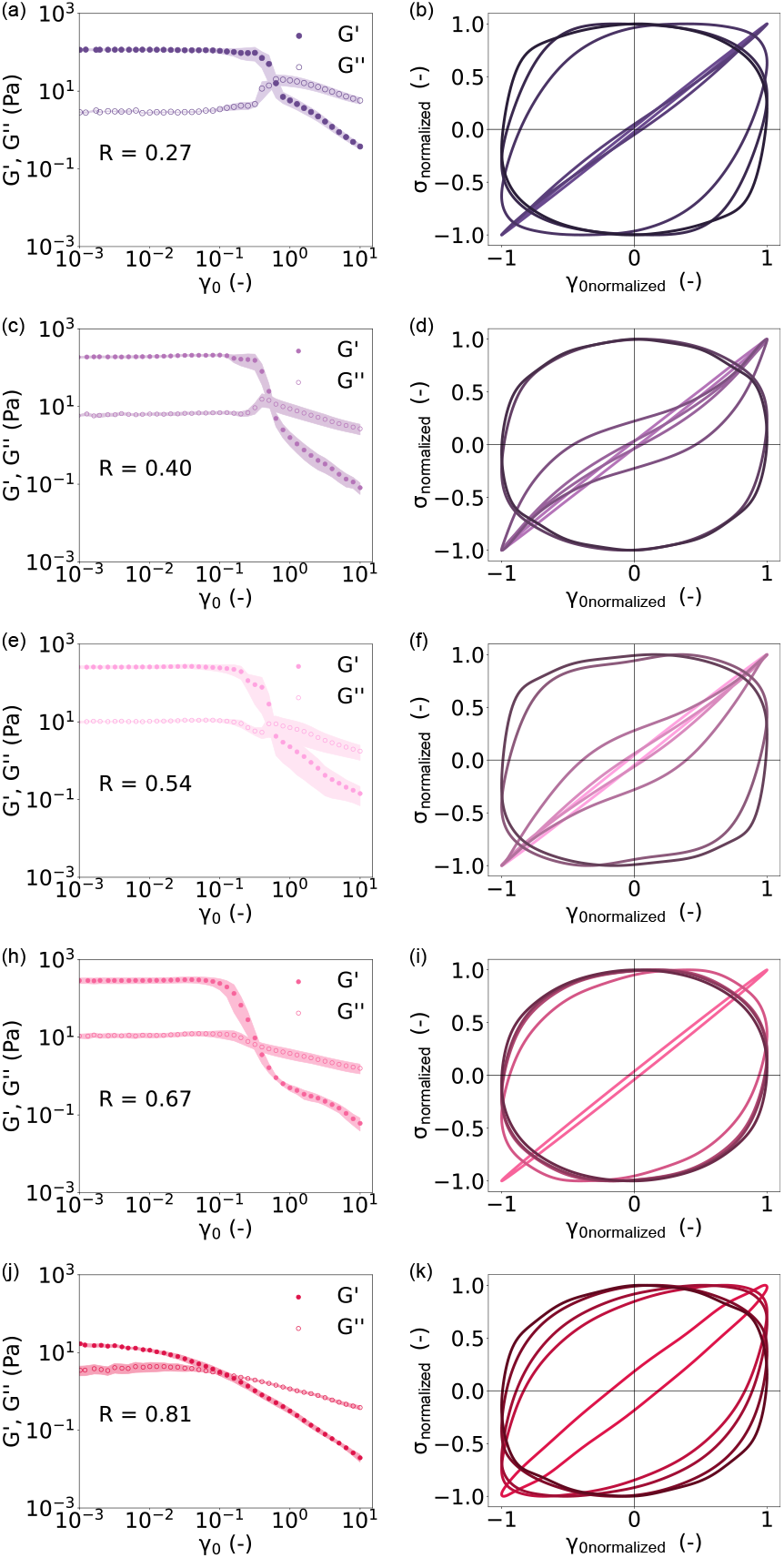
Rheological response to large amplitude oscillatory shear between (on the left) and within (on the right) the oscillatory cycles. Evolution of G’ and G” with increasing strain amplitude (*γ*_0_) of a hydrogel with *R* of (a) 0.27, (c) 0.40, (e) 0.54, (g) 0.67, and (i) 0.81. The shaded area represents the standard error of the measurements. Corresponding intracycle behavior of hydrogels is depicted by elastic Lissajous projections for different *R* values ((b) 0.27, (d) 0.40, (f) 0.54, (i) 0.67, and (k) 0.81 in five oscillatory cycles with *γ*_0_ *∼* 0.01, 0.25, 0.50, 1.60, and 6.30. The *γ*_0_ of oscillation increases from light to darker shades.

The differences in the dynamics of hydrogels below and above *R* ∼ 0.67 in Figure 2 and Figure 3(h) agree closely with the change in microstructure observed in fig. 1 (c). To examine this further, we examine the stress response (*σ*(t)) to the strain (*γ*(t)) in the cycle using elastic Lissajous-Bowditch (LB) projections. Corresponding to the average changes in G’ and G” with *γ*_0_, fig. 3 shows the elastic LB projections at different *R* for values of *γ*_0_, ranging from the LVE regime to the non-LVE regime for hydrogels. In the LVE regime, the stress response to a sinusoidal oscillatory strain is also sinusoidal, so at low *γ*_0_, the elastic LB projections take an elliptical shape as observed in Figure 3. As a hydrogel transitions to the non-LVE regime, *σ*(t) becomes distorted, (42) and the characteristic elliptical shape of the LB curves vanishes. This transition is observed for all samples at low *γ*_0_ but varies strikingly with *R*. At low *R* (*<* 0.40), the projections transition from a elastic to viscous-dominated regime at *γ*_0_ ∼ 0.5. This aligns closely with apparent the brittle yielding transition observed in Figure 3 (a) & (c) and likely indicates the dissociation of egg-box dimers. At high *R* (≥ 0.67), this transition is notably gradual and occurs at lower *γ*_0_. Interestingly, at intermediate values of *R* (0.40 - 0.54) and intermediate *γ*_0_ (∼ 0.5), a third type of response is observed, showing the emergence of distinct sigmoidal curves (Figure 3(d)-(f)). Such curves are often observed in biopolymer hydrogels (43–45) and are associated with strain-stiffening behavior. This strain stiffening is a general feature of semiflexible polymers, of which alginate is an example (46, 47), and arises from the pulling out of entropic undulations as the polymer chains are extended and stretched (48). As observed here, the onset of stiffening in alginate networks typically occurs at *γ*_0_ ∼0.5 (49) and is followed by the microstructural breakdown of the network at higher strains. Interestingly, we observe this strain stiffening only within a narrow regime of crosslinking stoichiometry (0.40 < *R* < 0.54). The prevalence of sigmoidal-shaped elastic LB projections can be visualized quantitatively by examining the third harmonic of stress response (*e*_3_) where positive values of *e*_3_ represents strain-stiffening (SI Appendix, Fig. S11). The sigmoidal curves at *γ* ∼ 0.5 transition into a curvilinear parallelogram waveform signifying microstructural breakdown, as *R* increases to 0.67 and beyond. The reasons for this are difficult to identify since the presence of strain stiffening is sensitive to multiple factors including (1) the polymer persistence length, which describes the bending rigidity of the polymer (50); (2) the crosslinking density, which defines how the undulations between crosslinking junctions are suppressed (51) and; (3) the microstructure of the network (52), which defines how the bulk shear strain maps onto the local strain. To rationalize the emergence of strain stiffening within this narrow parameter space, we must consider each of these factors in turn. For *R* < 0.5, an increase in *R* can be expected to initially increase the crosslinking density as more egg box dimers are formed, leading to denser crosslinking, which is expected to induce strain stiffening (50). For *R >* 0.5, the picture is more complex because higher order bundle-like structures are expected to form (6). A full theoretical analysis of the stiffening is beyond the scope of this work but briefly, the formation of bundles would be expected to increase persistence length and enhance strain stiffening so long as polymers within the bundle are strongly coupled (53). Previous x-ray analysis of alginate bundles suggest their lateral binding is somewhat disordered (21), which could imply the coupling between polymers is weak, which would prevent any expected enhancement in strain stiffening. Moreover, the formation of bundles could increase the hydrogel mesh size, resulting in a lower effective density of crosslinks, which would also be expected to actively suppress stiffening. We speculate, therefore, that the emergence of strain stiffening at *R >* 0.4 may arise from the increase in crosslinking density, while the disappearance of stiffening at *R* ≥ 0.67 may arise from the decrease in effective crosslinking density and weakening of lateral binding when bundle-like structures form. The observation that stiffening disappears precisely within the regime in which bundles are expected to form (6, 54) lends credence to this hypothesis.

This transition away from strain-stiffening behavior suggests that the hydrogels with *R* ≥ 0.67 have a reduced effect of rod-like rigid egg-box structure, highlighting a further structural evolution from clustered multiple egg-box strands that are prevalent at *R* ≥ 0.40. We surmise that this structural change related to the abundance of Ca^2+^ ions at high *R* can affect the molecular structure of crosslinked alginate through two modes. Firstly, higher Ca^2+^ concentrations screen the negative charge of the carboxyl (-COO^-^) units on the backbone of the alginate chains causing a reduction in repulsive forces and enabling lateral association of the crosslinked chains to form bundle-like structures as noted for analogous physically crosslinked networks (55). Secondly, higher Ca^2+^ concentrations decrease the bond life of the dynamic egg-box crosslinks making the chains in the bundles susceptible to deformations when strain is applied. Thus, the chains that form bundle-like structures can slide along the bundles dissipating energy easily at lower *γ*_0_ and thereby, increasing the viscous contributions. This is manifested in Figure 3(h) as the LVE regime shrinks considerably for *R* ≥ 0.67, and in Figure 3(i) with increased viscous contributions at *γ*_0_ ∼ 0.25 for hydrogels with *R* ≥ 0.67, which is much earlier than hydrogels with lower *R*. Morphologically, atomic force micrographs of alginate biopolymers with high R have also shown a larger apparent cross-sectional width (54) which can be surmised to be due to the formation of bundle-like structures.

### Bundle-like structures in hydrogels result in entirely distinct yielding behaviors

As a final test of the observed structural changes, we probe the hydrogels’ yielding dynamics under strain. Generally, investigating elastic stress (*σ*_*elastic*_) in a material is considered a useful estimate of yielding (56–58) and the approximate yield point is interpreted as the point at which the elastic stress component (*σ*_*elastic*_) reaches a maximum value and then subsequently decreases. Figure 4(a) depicts the differences in yielding behavior of hydrogels with different *R* values, characterized by the transition of *σ*_*elastic*_.

**Fig. 4.**
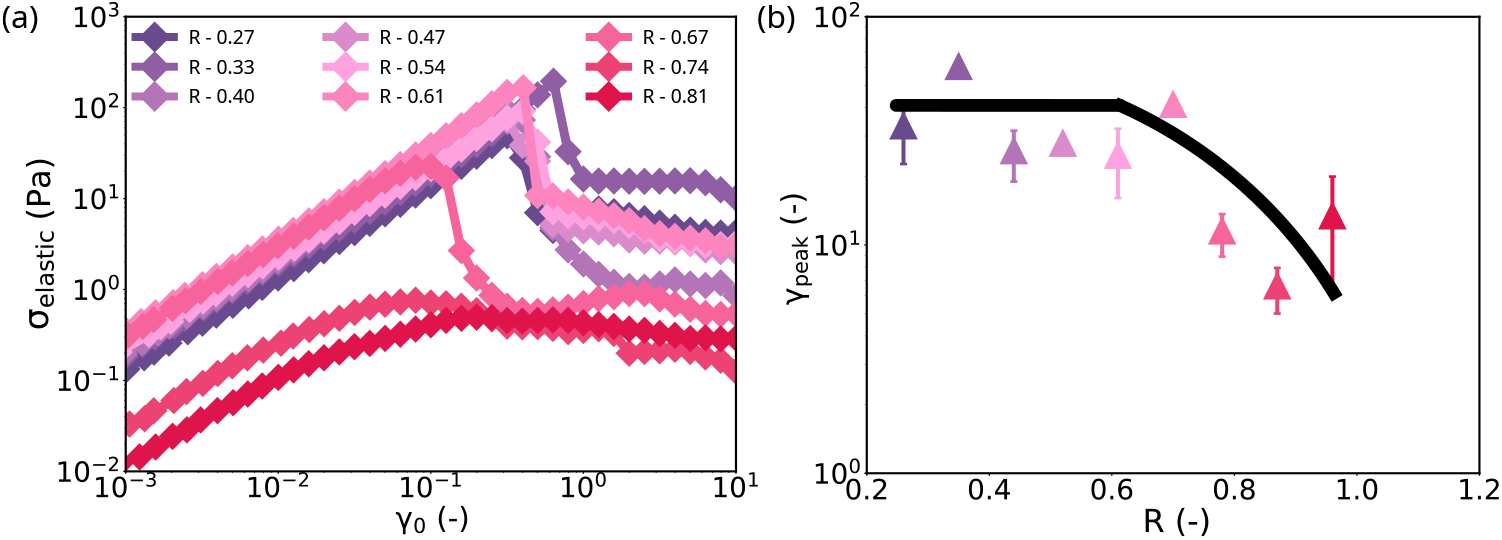
Characterization of yielding of alginate hydrogels through evaluation of the elastic contribution (*σ*_*elastic*_) to the total stress. (a) Evolution of *σ*_*elastic*_ with increasing *γ*_0_ for hydrogels with different *R* values. The peak in *σ*_*elastic*_ marks the yield point (b) Evolution of the average *γ* at the peak (*γ*_*peak*_) of *σ*_*elastic*_ in (a) with increasing Ca^2+^ concentration. The guide-to-eye solid line highlights the trend observed in *γ*_*peak*_.

Again, a clear transition is observed when *R* is increased: the transition in *σ*_*elastic*_ is relatively pronounced for *R* ≤ 0.67 and notably gradual and continuous for hydrogels with higher R. We explore the nature of microstructural changes directly by using a parameter known as the cage modulus (*G*_*cage*_) (SI Appendix, Fig. S12). *G*_*cage*_ describes the linear elasticity of the microstructure of the hydrogel network and is evaluated at the point in the rheological response of the hydrogel when there is a momentary stress equilibrium in each oscillatory cycle (59). This zero-stress point in the rheological response is indicative of the changes in the microstructure. At low *R* values, the normalized value of G_*cage*_ remains close to unity, aligning with the LVE regime of the hydrogels, suggesting no structural changes take place. When *γ*_0_ is increased beyond the LVE regime, *G*_*cage*_ shows a transition to lower values highlighting the structural breakdown of hydrogels. For hydrogels with *R* < 0.5, the final value of the normalized *G*_*cage*_ is approximately 60% of the initial value, suggesting only partial structural breakdown. As *R* increases above 0.5, this final value decreases to *<* 40%, indicating that an increase in *R* leads to a more complete structural breakdown. At highest *R* values (≥ 0.67), the evolution of *G*_*cage*_ is continuous, even at low strains with no apparent initial plateau in *G*_*cage*_. This suggests that hydrogels with high *R* values are highly susceptible to microstructural rearrangements even at low *γ*_0_. To summarize this, Fig. 4(b) presents the resulting yield point of the hydrogels in terms of the *γ*_0_ at which the first peak in *σ*_*elastic*_ is observed. We observe that for hydrogels with *R* ≤ 0.61, *γ*_0_ corresponding to the presence of the first peak in *σ*_*elastic*_ is similar. For higher *R*, the peak shifts to lower *γ*_0_. Our interpretation of this change in yielding is as follows. Hydrogels with high *R* possess bundle-like structures and these bundles are known to be amorphous and governed by weak noncovalent interactions .(21) Because of this lower bond-life time, the crosslinked alginate chains can readily remodel under shear. It is likely that this chain-sliding mechanism brings into effect a ductile yielding transition for hydrogels with high *R* as observed in Fig. 3.

## Discussion

We have carried out a cross-length scale characterization of the rheological and structural transitions of alginate hydrogels at a broad range of crosslinking stoichiometric ratios from 0.27 to 0.81, as informed by the extensive literature on alginate structure. We show an unambiguous transition from homogeneous networks to ordered bundle-like structures as confirmed by confocal microscopy. Aside from being structurally different, we show that the rheological properties of these hydrogels are also distinct. Counterintuitively, the microviscosity of bundle-like hydrogels appears to be significantly lower despite the fact that these hydrogels have higher crosslink densities. Furthermore, these trends clearly propagate to the macroscopic scale, with the bulk loss tangent increasing sharply around the transition to bundle-like structure. An examination of the nonlinear rheological properties of these hydrogels allows us to consolidate these two distinct length scales - both the observed structural changes at high *R* (Fig. 1(a)) and changes in microviscosity (Fig. 2(a)) correlate closely with a transition from brittle to ductile yielding, indicating an entirely distinct mechanism of microstructural breakdown under shear. By quantifying the nonlinear dynamics, the yield point and the cage modulus we suggest a perspective on how structure, rheology and yielding are all codependent on crosslink stoichiometry, summarized schematically in Fig. 5(a). Here, the left panel shows the commonly presented view of alginate hydrogel rheology: the structure is microscopically homogeneous, the crosslinking is governed by so-called egg-box dimers and the yielding mechanism is brittle in nature. The right panel summarizes the material characteristics of the bundle-like structures we observe and characterize in this study.

**Fig. 5.**
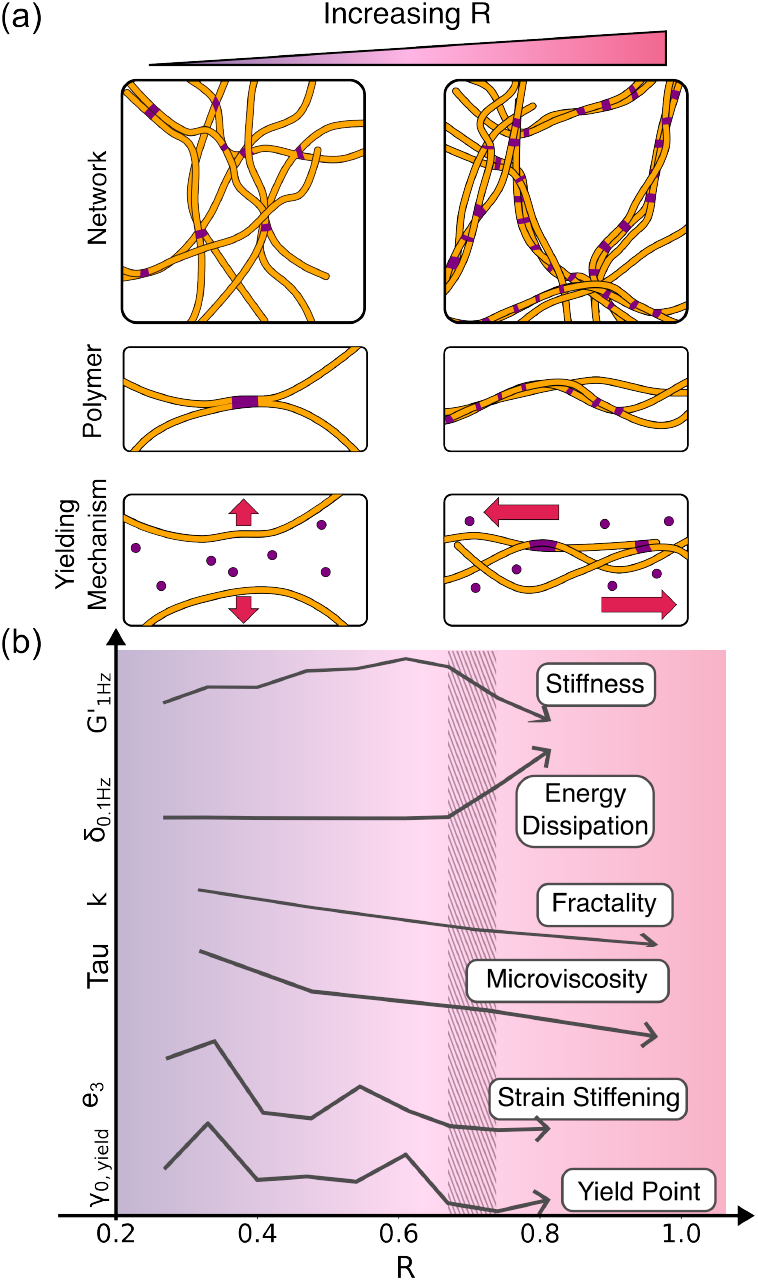
(a) Schematics highlighting the alginate network structure, crosslinking mode, and deformation mechanism at stochiometric ratios, R. At high R, (shaded region) networks become coarse due to the formation of bundle-like structures. The presence of these structures also causes the transition from a brittle-like to a ductile-like yielding regime. This is supported by six key metrics that are quantified in this study (b) relating to rheology, microviscosity, structure and yielding.

Figure 5(b) presents the transition between these two states, plotting each of the structural and rheological parameters examined here with respect to R. By consolidating these metrics, we identify the key trends associated with the transition from network-like to bundle-like structures, which appears to occur for 0.67 *> R*. When considered from the perspective of linear viscoelasticity, this transition appears relatively sharp because the values of G’ and *δ* decrease unambiguously after this point. However, our microscopic characterization presents a different picture. The value of k, which describes network fractality, decreases steadily across all values of R, even those far from the transition point, indicating a continuous structural change. Similarly, the value of Tau, which reports the microviscosity of the sample, also decreases steadily across all values of R. Remarkably, the nonlinear rheological properties, manifested through the third nonlinear harmonic *e*_3_ and the yield point *γ*_0,*yield*_, also decrease gradually across all *R* values. This strongly suggests that the observed transition from brittle to ductile yielding is associated with these microstructural changes. A possible mechanistic origin for this behavior can be inferred from the known physical properties of alginate’s bundle-like form. Small angle scattering (21) of alginate in solution has shown that these bundle-like structures, unlike the egg-box dimers found in the network-like structures, have amorphous packing governed by noncovalent interactions. It is likely that these associations between alginate chains in the bundle-like form are weaker than those in the network-like form, which would clearly explain why the yielding process that we observe is ductile and continuous.

## Conclusions

By combining imaging approaches, fluorescence lifetime imaging and rheological approaches we are able to correlate the structure and rheology of alginate hydrogels across a broad range of crosslink stoichiometries. We identify the emergence of a bundle-like structure, at high values of the stoichiometric ratio, *R*, which has previously only been observed in alginate solutions. Through a multi-modal approach we show that these bundle-like structures exhibit a coarser microstructure and possess a reduced microviscosity, a lower elastic modulus and a higher degree of energy dissipation. The resulting bundle-like hydrogels undergo yielding in a way that is fundamentally different to the network-like structures formed at low crosslinking stoichiometry. It seems likely that this distinct yielding mechanism results from the amorphous cross-sectional structure of the alginate bundles since the loose disordered bonds within the bundle may lead to a more compliant rheological response. This would lead to a gradual process of microstructural breakdown with strain which is consistent with our observation that the networks are highly sensitive to shear strain (Figure 4) and the gradual reduction in the apparent G’ (Figure 3).

The ability to predictably control the viscoelasticity of alginate hydrogels is an integral ambition in a wide variety of applications including food, packaging and biomedical devices. In all of these, a common strategy is to tune the viscoelasticity by adjusting the crosslink stochiometry(60). Here, we show that the reality is more nuanced because, at high R, increasing the crosslinking in fact decreases the elastic modulus. Our observation that, in this case, *less is more* has fundamental implications for the formulation of alginate-based products. Furthermore, alginate biopolymers are known to adopt different structures within the living cell wall (18), with thick bundle-like structures co-localising with regions that are subject to high wall stress. It will be interesting to examine whether the regulation of alginate structure that we observe here *in vitro* turns out to be involved in stress responses in living cell walls.

## Materials and Methods

Our experiments used DuPont Protanal CR 8133 sodium alginate powder, which has a molecular weight of 135 kDa. Calcium carbonate (CaCO_3_), and glucono-*δ*-lactone (GDL) were purchased from Sigma Aldrich.

To carry out optical imaging studies, rhodamine B (RhB) from Acros (now Thermo Scientific Chemicals, part number 13211000), RhB-labelled alginate from Creative PEGWorks (AL510, low viscosity 100-300 cP, 1% mol, SI Appendix, Fig. S1) and glycerol of highest purity (Avantor, 32450.AE) were used without purification. Alcohols — including methanol, ethanol, 2-propanol, n-butanol, and tert-butanol, were of chromatographic purity and used as received.

### Characterization of alginates using 1H Nuclear Magnetic Resonance (NMR)

To characterize M:G ratio of alginates using NMR spectroscopy, a preliminary partial hydrolysis step to reduce the molecular weight of the polymer was performed. This process decreases the solution viscosity and allows the acquisition of spectra with well-resolved signals. The process of hydrolysis was carefully controlled to avoid complete polymer degradation or epimerization of G and M monomers, since this could interfere with M:G ratio measurements. Solutions of alginate powder and deionised water (1 mg/mL) were prepared under continuous stirring until a homogeneous sample was obtained. The pH was adjusted to 5.6 using HCl to start the hydrolysis, stirring continuously at 100°C for 1 hr. After cooling the sample to room temperature, the pH was adjusted to 3.8 and stirred for 30 mins more, at 100°C. After cooling the sample, the solution was neutralised by adjusting the pH to 7-8 using NaOH. The samples were freeze-dried overnight, and then redissolved in D_2_O and freeze-dried again to remove residual water. After the second step of freeze-drying, the samples were redissolved in 1mL of D_2_O; 700 L of this solution was transferred into 5 mm NMR tubes.

The spectrometer probe was stabilised at 80°C, and the parameters used were: Spectral width: 15 ppm, Acquisition time: 4 s, Relaxation delay: 2 s, and Number of scans: 64–128. The NMR data processing includes Fourier transformation without zero filling, manual phase and baseline correction, spectral shift using solvent (D_2_O) as a reference, manual detection of peaks for region B, and deconvolution. Region B allows for estimating the sequential distribution of residues (diads and triads), providing information about MM, GG, and MG blocks along the alginate chain. The alginate composition is determined by integrating the signals corresponding to the anomeric protons and H-5 protons of the M and G groups. For DuPont CR8133 protanal, M:G ratio was found to be 1.23 (SI Appendix,Fig S13) while for AL510, the ratio is 1.43 (SI Appendix,Fig S14).

### Hydrogel preparation

To construct hydrogels, a stock solution of 0.4 mM alginate was prepared in Milli-Q water and stirred for at least 15 minutes at 90^*°*^C until fully dissolved. Stock solutions of various CaCO_3_ concentrations were prepared by mixing the appropriate mass in Milli-Q water, followed by sonication for 20 minutes. The stock solution of GDL was prepared by vortexing the required mass in distilled water for 20 seconds. Reaction mixtures of 2.5 ml were made by adding different amounts of alginate, CaCO_3_, GDL, and Milli-Q water while giving a final alginate concentration of 0.12 mM. We define *R* as charge equivalency ratio (= 2*[Ca^2+^]/[G units]) (61). The *R* value of a hydrogel sample determined the concentration of CaCO_3_ and the amount of GDL was then adjusted to keep a consistent GDL: CaCO_3_ molar ratio of 0.18. Since alginate can convert to alginic acid at pH 3.6, the pH of the hydrogels was monitored during formation and was found to stabilize between 5.5 and 7.5 on gelation (SI Appendix, Fig. S2). For microscopy experiments, a home-made chambered microscopy slide, which consisted of the glass coverslip (22×50 mm, Class 1.0 borosilicate glass, VWR) and a plastic grid from an 8-well chambered microscope slide (Lab-Tek Nunc II, part number 155411, ThermoFisher Scientific, Germany) mounted on the glass coverslip and fixed with two tiny drops of superglue was used. The hydrogels were prepared according to the above protocol with the only change in the total volume, which was 50-70 L. Hydrogel components (alginate, CaCO_3_ and GDL) were mixed in proportions required for each *R* value (0.29, 0.44, 0.65 and 0.87) and deposited into individual wells for gel formation for two hours. To prevent sample drying, the whole slide was sealed with another coverslip and parafilm.

### Rheology

The rheometer used for all oscillatory tests was a Netzsch Instruments Ltd. Kinexus Pro+. All experiments were carried out at a controlled temperature of 25^*°*^C using a 40 mm, cone plate upper geometry with a 1^*°*^ angle. Before loading samples the inertia of the cone geometry was calibrated and a rotational torque mapping program was carried out to determine the angular position of the geometry at which the torque variation was the lowest. The decimator of the rheometer controller, a proprietary settings that controls the torque sensitivity, was manually adjusted to 5, which represents the highest position sensitivity attainable.

Once mixed, the reaction mixture was loaded quickly on to the rheometer lower plate with a pipette, and the upper geometry was lowered. Silicone oil was distributed on the lower plate around the edges of the upper geometry to prevent sample evaporation. The reaction mixture was pre-sheared at a rate of 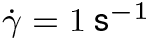 for 100 seconds to distribute the sample evenly between the plates and remove any deformation history caused by loading. The hydrogel gelation kinetics and linear viscoelastic properties were measured by carrying out small amplitude oscillatory shear rheology. To capture the hydrogels’ gelation kinetics, continuous measurements were carried out a frequency of 6.28 rad.s^−1^ and strain amplitude (*γ*_0_) of 0.005 for 1.5 hours. Next, a frequency sweep was carried out to measure the viscoelastic moduli, within the range of 628 to 0.0628 rad.s^−1^ at a constant amplitude of 0.005, which was within the LVE regime for all samples. Finally, to investigate rheological behavior in the non-LVE regime, LAOS waveforms were applied to examine how the sample responded under increasing strain. *γ*_0_ was increased from 10^−4^ to 10 at a constant frequency of 3.14 rad.s^−1^, imparting 10 cycles of oscillations at each value of *γ*_0_ to ensure steady state. Waveform analysis of the resulting Lissajous-Bowditch curves (stress vs. strain) was carried out on raw data of the final cycle. All rheological measurements were carried out in triplicate on independent samples.

### Absorption and Fluorescence Spectra, Fluorescence Lifetime Measurements

Rectangular quartz cuvettes (10 x 10 mm^2^, total volume 4 mL) were used for all spectroscopic measurements with RhB and AL510. Absorption and fluorescence spectra were recorded using an Agilent 8453 UVvis spectrophotometer and a Fluoromax-4 spectrofluorometer (Jobin-Yvon, Horiba), respectively. All liquid samples had an absorbance below 0.15 at the absorption maxima. All fluorescence spectra were corrected for wavelength-dependent sensitivity of detection. Time-resolved fluorescence decays were measured using a DeltaFlex setup (Jobin-Yvon, Horiba) operating via Time Correlated Single-Photon Counting (TCSPC). The excitation source was 467 nm NanoLED (Horiba, FWHM < 200 ps); the detection wavelength and spectral width were selected with the in-built monochromator as indicated. The fluorescence decays were recorded until 10,000 counts at the maximum and in 100 ns time window with 4096-time bins. Fluorescence decays were fitted globally using a sum of exponential decay functions (details are specified in the text). The resulting goodness-of-fit parameter *χ*^2^ was between 1.0 and 1.3. The instrument response function (IRF) was measured using a dilute Ludox solution at the excitation wavelength.

### Two-photon microscopy

Emission images were recorded using an inverted confocal laser-scanning microscope, Leica TCS SP5-II (Leica Microsystems Ltd, Germany) at room temperature. RhB and AL510 emission in the range of 540 - 680 nm was collected following two-photon 930 nm excitation from a Chameleon Vision-II Ti:Sapphire laser (Coherent Inc., Germany). A 20x objective (HC PL APO CS2, N. A. = 0.75, dry objective, part number 11506517, Leica) was used to collect images at 1024 x 1024-pixel resolution. All images were processed in the Leica Application Suite X (LAS X) software package.

### Fluorescence Lifetime Imaging Microscopy (FLIM)

Two-photon excited FLIM imaging was performed using the same microscope system described in the previous section and the same excitation and detection wavelengths. FLIM data were acquired with an SPC-830 photon counting card and Spcm64 software (both from Becker & Hickl). The resolution of FLIM images was 256×256 pixels with 256-time channels in pixel decays. The IRF was obtained by measuring the second harmonic generation (SHG) signal from crystals of urea. The FLIM data was analysed using the SPCImage software (Becker & Hickl, v.8.5) using 3 x 3 square binning (bin 1) and fitting with a monoexponential function.

## Data, Materials, and Software Availability

Raw rheological data, fluorescence and fluorescence lifetime images and computed fractality parameters will be available at the University of Liverpool Repository.

## ACKNOWLEDGMENTS

V. K., P.S., and S.G. are supported by LILACS project, funded by UK Research and Innovation (MR/Z505407/1). V.K. was also supported by an Engineering and Physical Sciences Research Council (EPSRC) PhD studentship through the Soft Matter for Formulation and Industrial Innovation Centre for Doctoral Training (SOFI2 CDT) (EP/L015536/1), co-funded by Reckitt, UK. RGG is supported by a PhD studentship, funded by the Fundación Politécnico in Mexico.

