## Supplementary Information for "Correlating Structure and Rheology in Ionically Crosslinked Alginate Biopolymer Hydrogels - A Case for Why “Less” can be “More”"

For

### Section: Chemical structure of RhB and RhB-labelled alginate and RhB-labelled alginate hydrogels

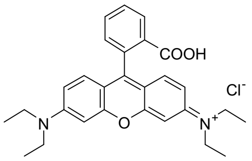


Rhodamine B

(RhB)

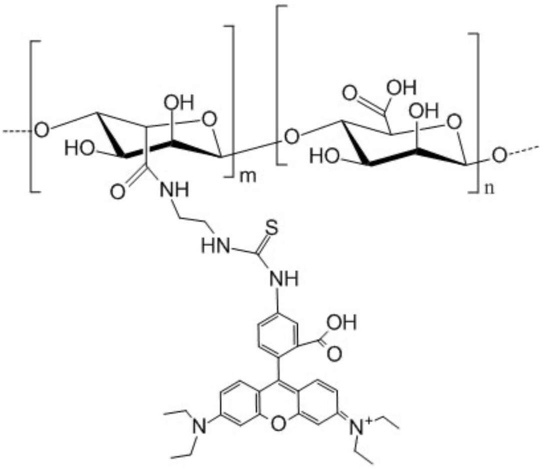


Rhodamine B-labelled alginate

(AL510)

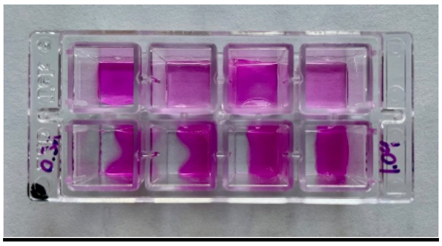

Figure S1: Chemical structures of rhodamine B (RhB) and RhB-labelled alginate (AL510). The structure for AL510 was provided by Creative PEGWorks. The hydrogels formed using RhB-labelled alginate at R=0.32, 0.48, 0.73, 0.97 from left to right.

(https://creativepegworks.com/product/alginate-rhodamine-low-viscosity-100-300-cp).

### Section: pH measurements during gelation

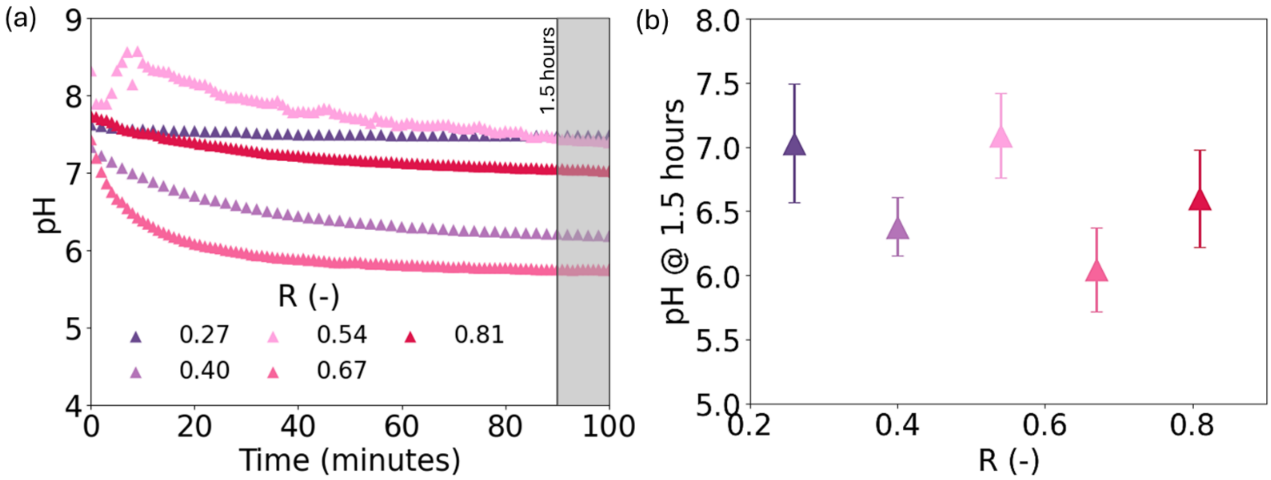

Figure S2: (a) Evolution of pH during gelation of alginate hydrogels for different R values; (b) pH on gelation after 1.5 hours for the hydrogels with different R values.

### Section: Fractal dimensions

Fractal dimensional analysis of the 2D image of the alginate gel network was calculated as described in (reference: https://pubs.rsc.org/en/content/articlepdf/2023/sm/d3sm00111c) using the Hausdorff–Besicovitch, or box-covering, dimension. In this method the system is divided into a series of sets of unit squares of width *r* starting with *r=system width* and halving *r* every time until a value *of r=pixel size* (ie *r=1* if *r* is in units of pixels) is reached. For each value of *r* the occupancy number *O(r)* is calculated as the number of boxes which contain at least one pixel above a specified threshold, in this case 50.

For a homogeneous bright system *O(r)* will simply equal the number of boxes at a given r, however if not all pixels are filled the relationship is non-linear.

For an ideal homogeneous network where the probability of any given pixel being unfilled is equal for all pixels *O(r)* will follow the equation:

$$O(r) = \frac{system\_width^2}{r^2}\left(1 - p^{r^k}\right)$$

Where p is the probability of any pixel being unfilled in the whole image and k=2

For a non-homogenous system however, the value of *k* will vary between 0 and 2, by fitting the above equation to a plot of *r* vs *O(r),* we can identify the value of *k* for each image.

Example fit: this is for the final *R= 0.97* image and gives *k=1.63±0.01*

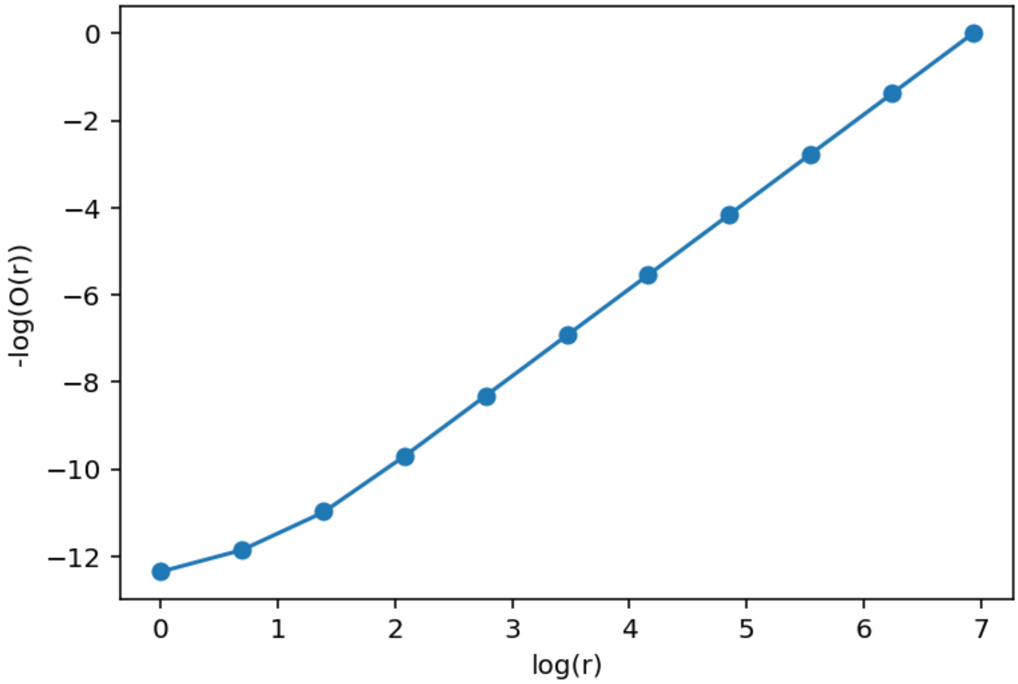

Figure S3: *O(r)* vs *r* dependence calculated for one of the images obtained with gel R = 1.04.

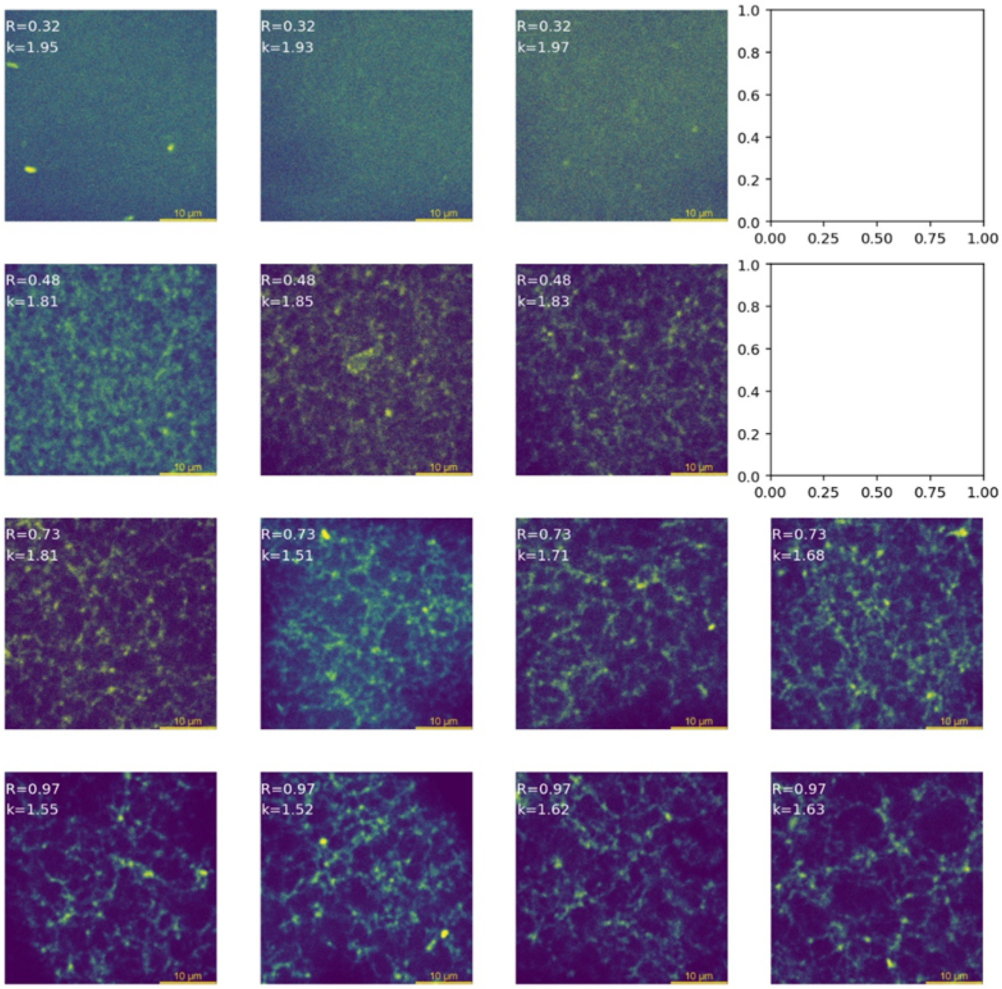


Figure S4: Confocal images of RhB-labelled alginate gels at different R values presented for the threshold of 150. The corresponding *k* values are given for each image.

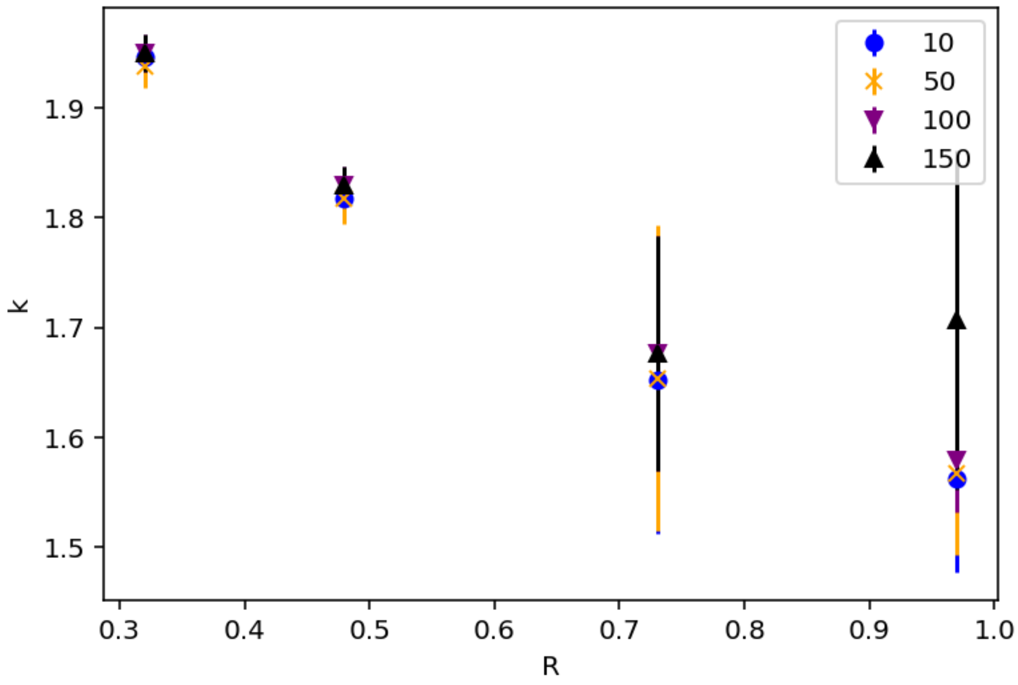


Figure S5: The *k* vs R plots for different threshold values in the analysis of confocal images in Figure SF2. The changes are only seen at above 150.

### Section: Fluorescence intensity and FLIM measurements

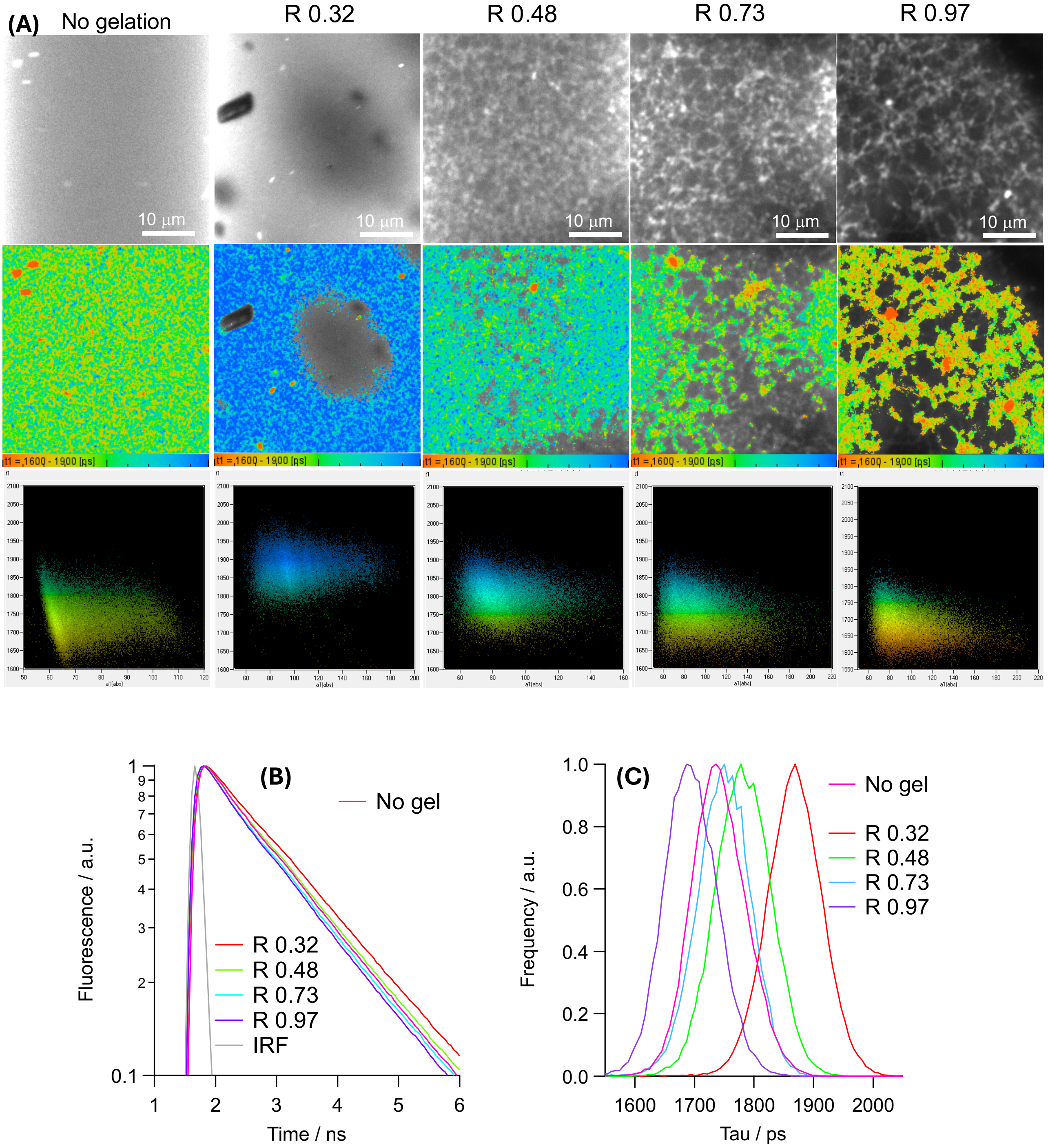


Figure S6: (A) Fluorescence intensity (upper row) and FLIM (middle row) images obtained with AL510 in aqueous solution (54 mg/ml) and gels (13.5 mg/ml, four R values) recording the emission in the 540-680 nm range following the two-photon excitation at 930 nm. Scale bars are 10 µm. (Bottom row) Correlations between intensity and lifetime for FLIM data in the middle row. (B) Decay traces averaged over the frame and (C) lifetime distributions for FLIM data in (C).

### Section: Study with binary mixture of water and glycerol

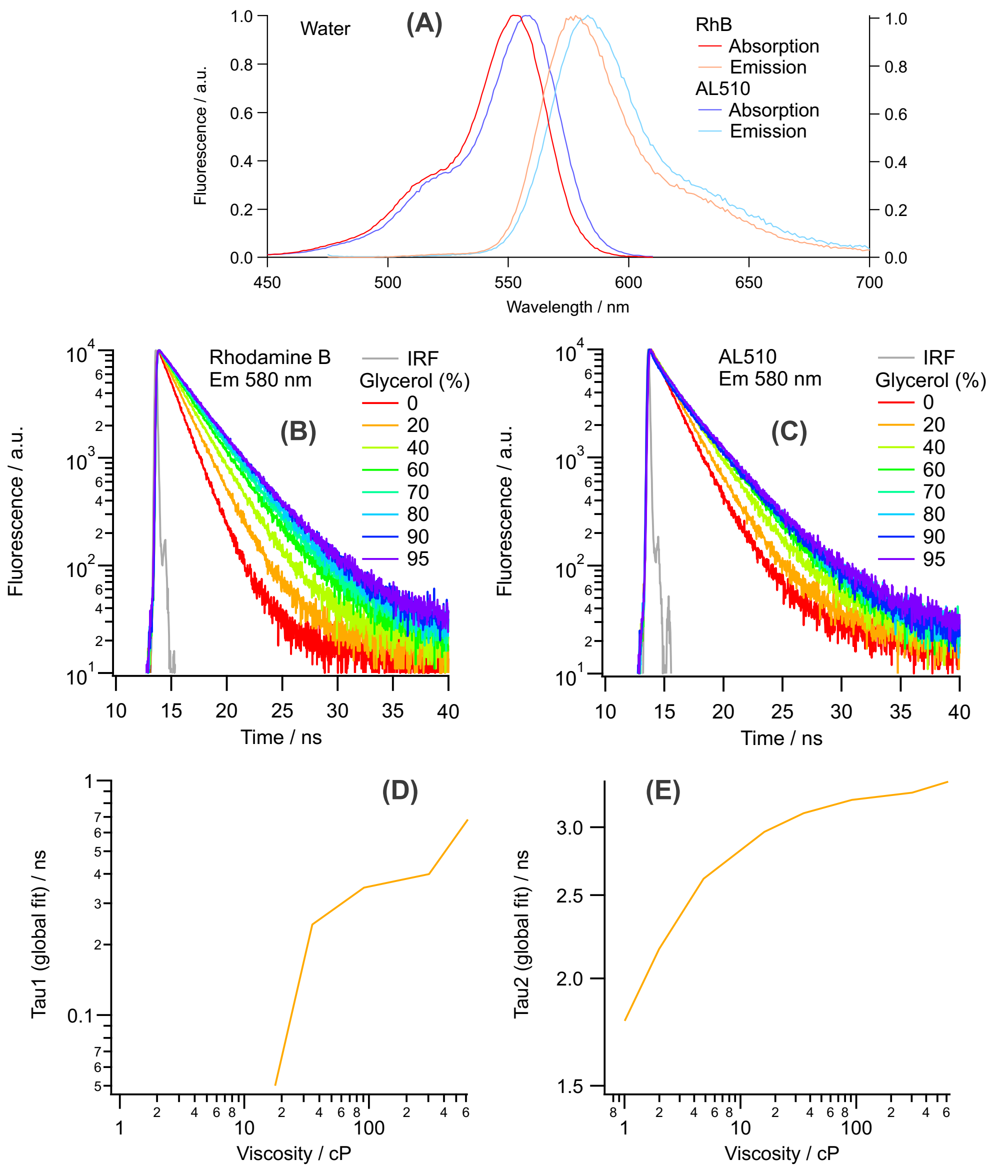


Figure S7: (A) Fluorescence emission and excitation (denoted as absorption) spectra of RhB and AL510 (both 0.15 µg/ml) in an aqueous environment. (B,C) Fluorescence decay traces recorded with (B) RhB and (C) AL510 at 580 nm after the excitation at 467 nm in binary mixtures of water and glycerol (annotations: glycerol content). (D,E) Viscosity calibrations for tau1 (solvent relaxation) and tau2 (fluorescence decay) obtained in the global analysis of decay traces recorded over the whole emission band (see Figure S8 for decay traces).

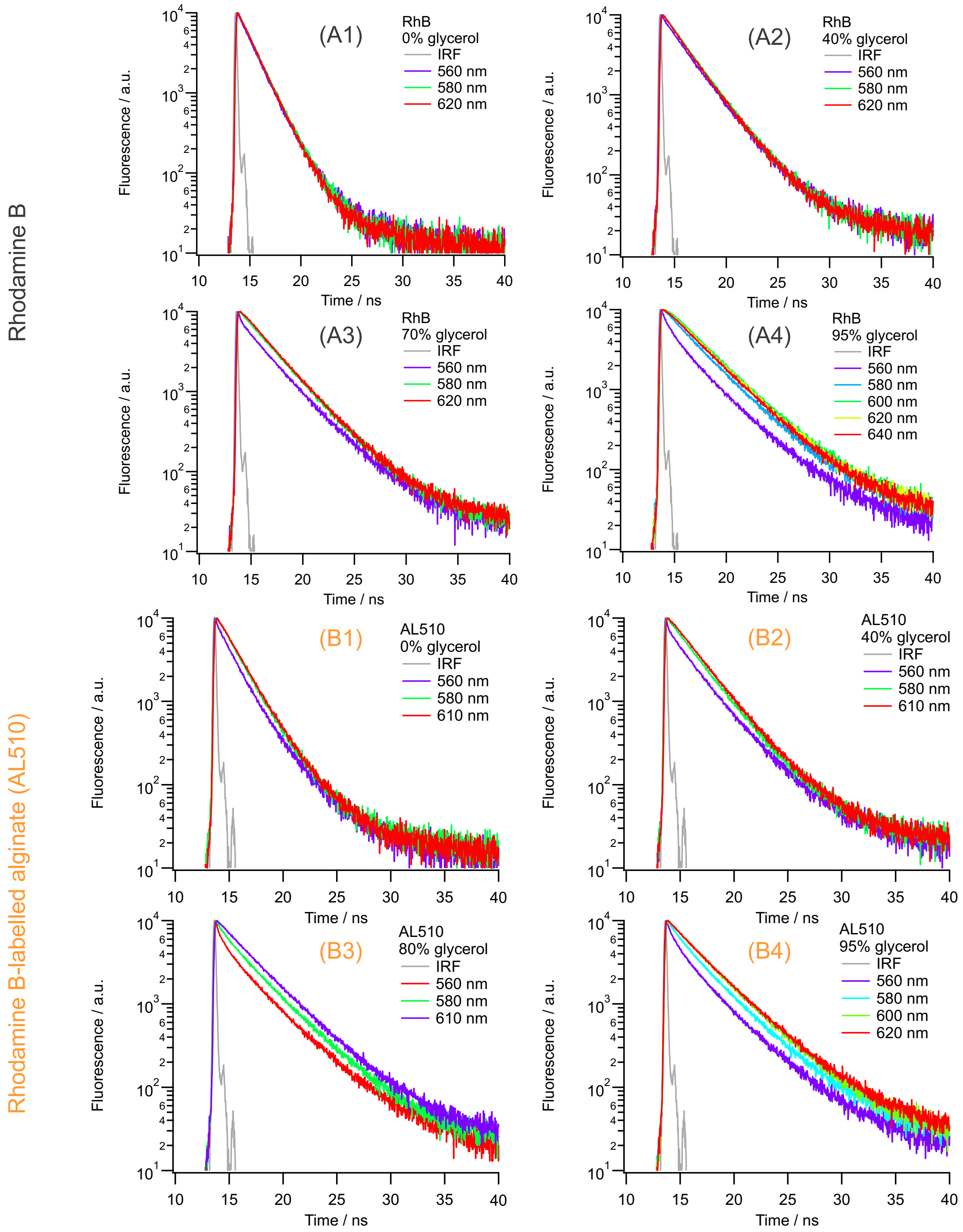


Figure S8: Fluorescence decay traces recorded for RhB and AL510 (both 0.15 μg/ml) over the whole emission band after the excitation at 467 nm in binary mixtures of water and glycerol. (Ai, i=1,..,4) RhB and (Bi, i=1,..,4) AL510 for (A1, B1) 0%, (A2, B2) 40%, (A3, B3) 70% and (A4, B4) 95% of glycerol in the mixture (by volume).

Data shown in Figure S8 were fitted in the global analysis using a sum of two or three exponential functions. Two-exponential model was used in all cases with coincident decay traces over the emission band, and the three-exponential model was used for all other cases to account for the solvent relaxation in mixtures of high viscosities. In all cases, the last component was used to reproduce long-lived decays (lifetimes > 30 ns) originating from an impurity present in all rhodamine

B samples (both RhB in the free state and bound to alginate). Due to a low amplitude (< 1%), this component was omitted from further considerations. The time resolution of the available TCSPC unit was considered as 100 ps. Therefore, all characteristic times below the time resolution limit were considered as values shorter than 100 ps.

Table S1. Characteristic times obtained from the global analysis of fluorescence decay traces recorded with RhB and AL510 in binary mixtures of water and glycerol at 20°C. Tau1 is considered the relaxation time, and tau2 is the fluorescence lifetime of RhB in the free state or bound to alginate (AL510). Viscosity values were calculated according to the following ref [https://www.met.reading.ac.uk/~sws04cdw/viscosity_calc.html].

| | | RhB | | AL510 | |
|---|---|---|---|---|---|
| Glycerol (v, %) | Viscosity / cP | Tau1 / ns | Tau2 / ns | Tau1 / ns | Tau2 / ns |
| 0 | 1.005 | – | 1.568 | < 100 ps | 1.785 |
| 20 | 1.985 | – | 2.008 | < 100 ps | 2.164 |
| 40 | 4.81 | – | 2.399 | < 100 ps | 2.613 |
| 60 | 16.1 | < 100 ps | 2.763 | < 100 ps | 2.963 |
| 70 | 35.1 | < 100 ps | 2.948 | 0.243 | 3.116 |
| 80 | 91.5 | 0.148 | 3.123 | 0.349 | 3.227 |
| 90 | 302.7 | 0.306 | 3.308 | 0.399 | 3.291 |
| 95 | 621.2 | 0.458 | 3.373 | 0.681 | 3.389 |

### Section: Control experiments with alcohols

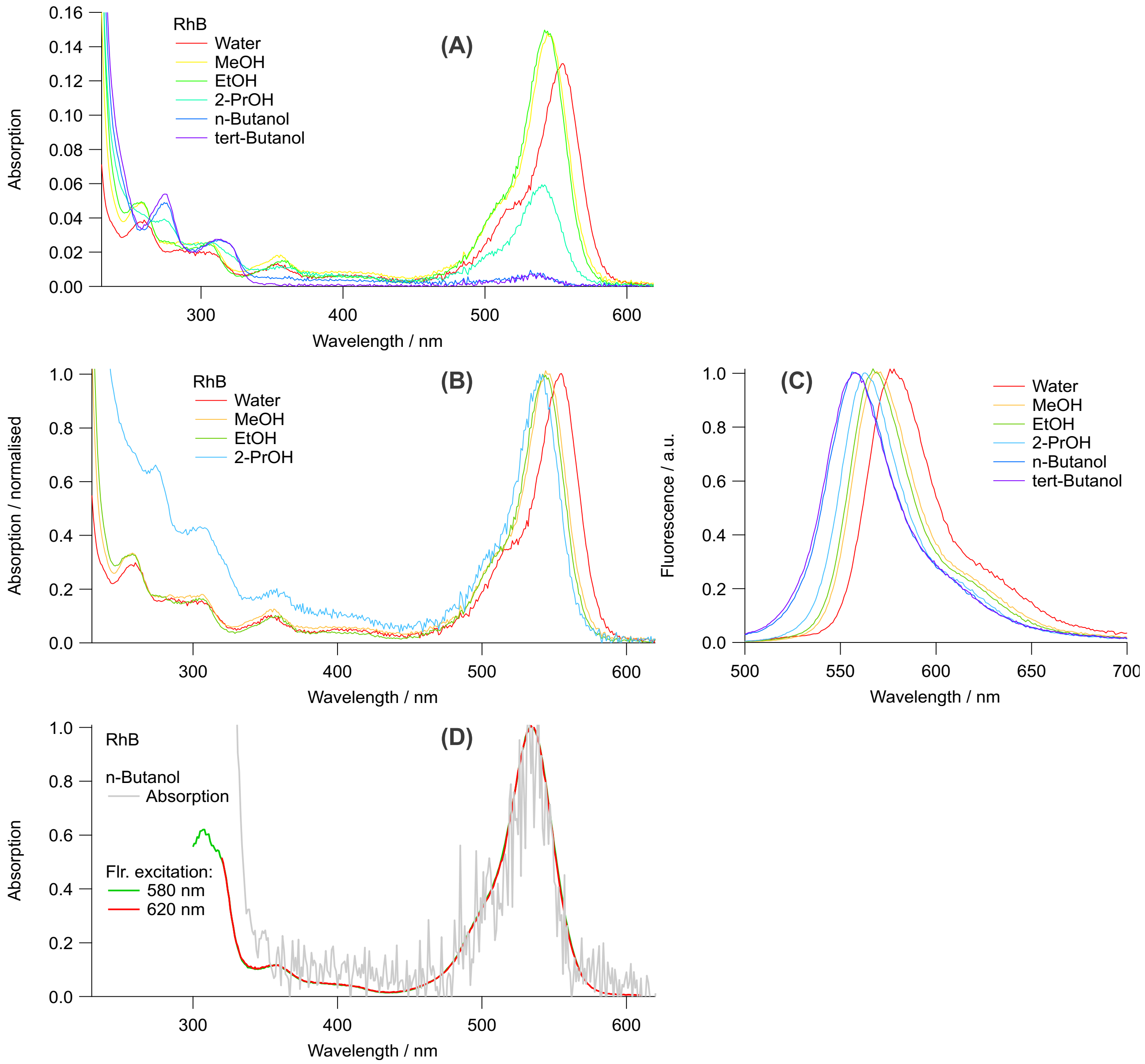


Figure S9: (A, B) Absorption and (C) fluorescence spectra for RhB in several alcohols. Fluorescence spectra were recorded after excitation at 490 nm. (D) Comparison of absorption and fluorescence excitation spectra for RhB in n-butanol.

RhB is low soluble in n-butanol and tert-butanol, but it exhibits measurable fluorescence, see Figure S9(C). Complete agreement between the absorption and fluorescence excitation spectra (Figure S9(D)) confirms that the observed fluorescence spectrum (Figure S9(C)) originates from RhB in n-butanol and tert-butanol.

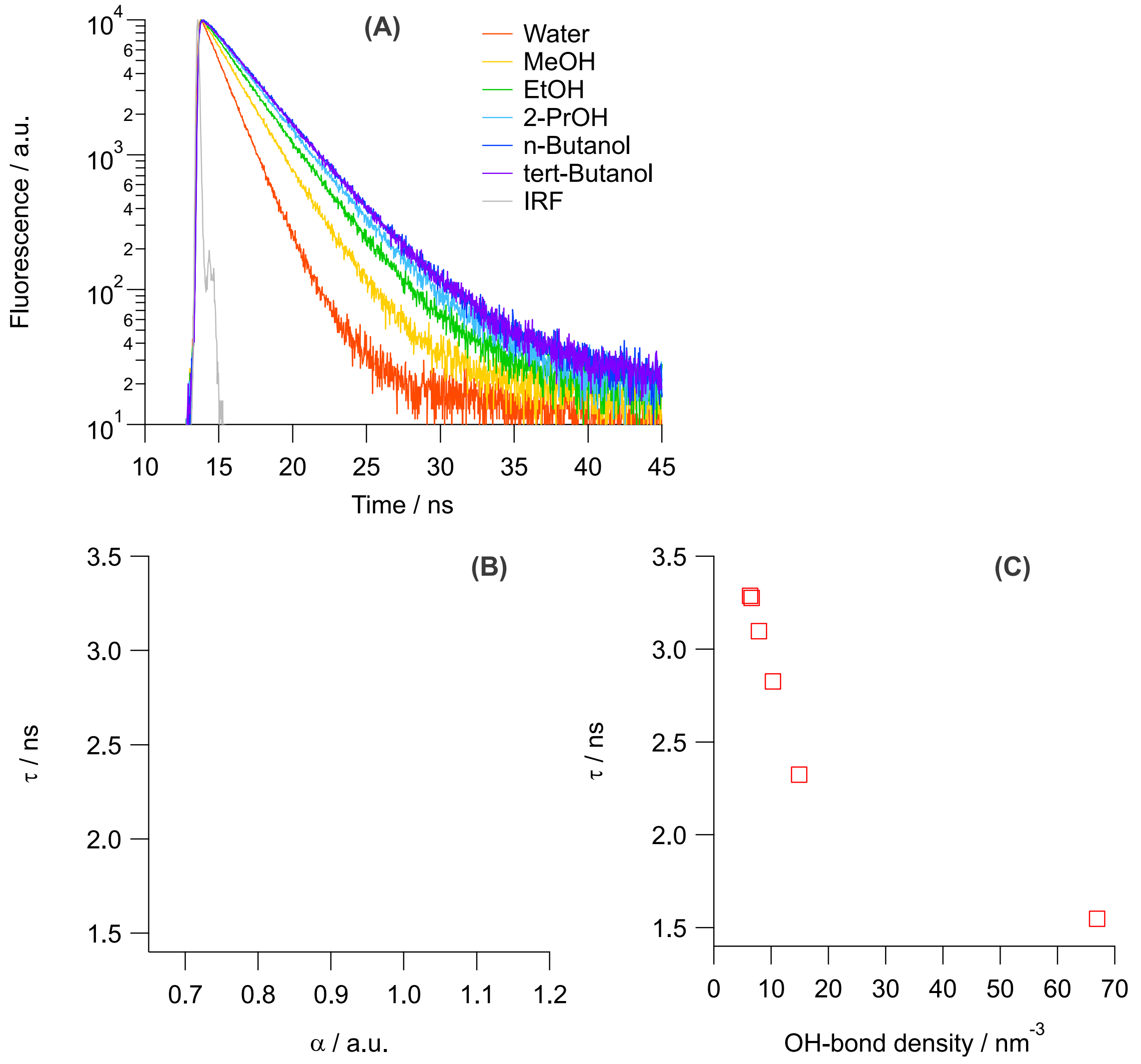


Figure S10: (A) Fluorescence decay traces recorded for RhB (0.15 µg/ml or 0.3 µM) in different alcohols at 580 nm after the excitation at 467 nm. (B,C) Correlation between the lifetime of RhB in different alcohols and (B) the alpha parameter of Kamlet-Taft's scales [2] describing the solvent's ability to donate the hydrogen bond, and (C) the density of OH bonds calculated as the inverse of volume occupied by a single solvent molecule using the molecular weight and the solvent density.

Table S2. Solvent dependence of $\alpha$ parameter describing the H-bond donation ability, density of OH bonds and the fluorescence lifetime of RhB.

| Solvent | $\alpha$ [2] | $\rho_{OH}$ / nm$^{-3}$ | $\tau$ / ns |
|---|---|---|---|
| tert-Butanol | 0.68 | 6.35 | 3.286 |
| n-Butanol | 0.79 | 6.58 | 3.274 |
| 2-Propanol | 0.76 | 7.88 | 3.096 |
| Ethanol | 0.83 | 10.31 | 2.826 |
| Methanol | 0.93 | 14.89 | 2.323 |
| Water | 1.17 | 66.91 | 1.548 |

**Section: Third harmonic of stress response**

To characterise the intracycle behaviour quantitatively, the hydrogel response was decomposed using Chebyshev polynomials [3] and examine the elastic Chebyshev coefficients present in the stress response: $e_1$, and $e_3$. $e_1$ in Figure S11(a) represents the first Chebyshev coefficient in applied strain within an oscillatory cycle that correlates to the linear behaviour of the hydrogels, whereas $e_3$ in Figure S11(b) represents the third Chebyshev coefficient and represents the mathematical criteria for specifying the physical interpretation of the nonlinearity such as strain-stiffening behaviour. When the sample exhibits strain-stiffening behaviour under an oscillatory cycle, $e_3$ is positive, whereas a negative $e_3$ highlights the strain-softening behaviour. As observed in Figure S11(a), $e_1$ for all hydrogels remains constant at low strain amplitude ($\gamma_0$) highlighting the LVE regime of the hydrogels. As $\gamma_0$ increases beyond the LVE regime of the hydrogels, $e_1$ decreases for all the hydrogels as higher terms emerge in Chebyshev decomposition of the stress. $e_3$ is near-zero in magnitude at low $\gamma_0$ in the LVE regime and at very high $\gamma_0$ (>= 1) when higher terms are present. However, we observe a spike in $e_3$ for the intermediate $\gamma_0$ between 0.1 and 1 at R <= 0.67. This region represents the $\gamma_0$ and R where significant strain-stiffening is observed and corresponds to the prevalence of sigmoidal-shaped elastic LB projections. We note that the ratio of $e_3$ to $e_1$, $e_{3/1}$, known as the relative third harmonic, is presented as an alternative measure of strain stiffening [4-5]. In this case, $e_{3/1}$ is near-zero for $\gamma_0$ <= 0.1 and is only observed to increase beyond $\gamma_0$ ~ 0.1 irrespective of R values (Figure S11(c)). We attribute this distinction in features of $e_3$ and $e_{3/1}$ to the sharp decline in $e_1$ at high $\gamma_0$ as $e_1$'s decline is observed to increase $e_{3/1}$ and as a result, $e_{3/1}$ might not be an adequate parameter to highlight the strain-stiffening behaviour of the hydrogels.

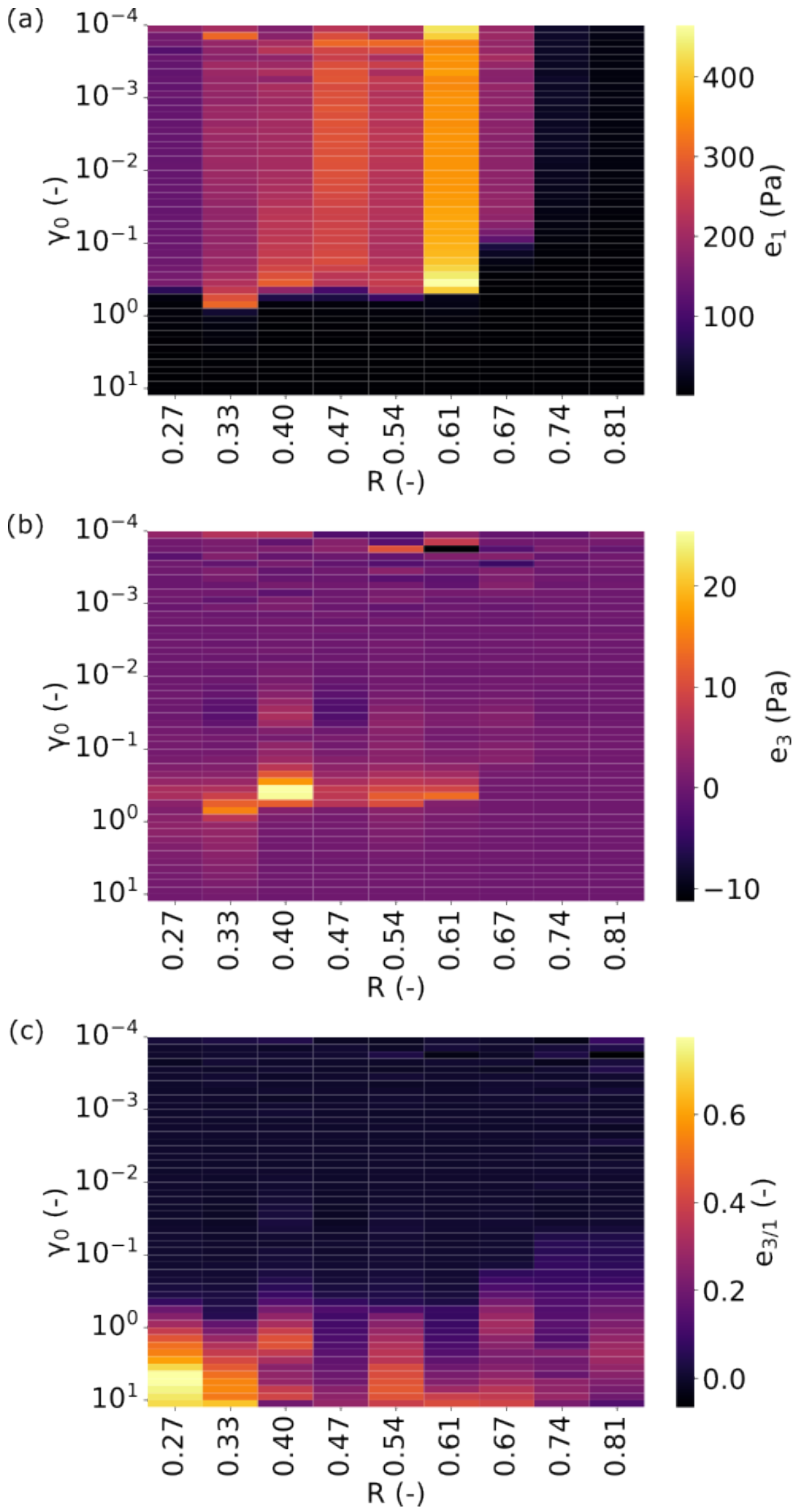


Figure S11: Harmonics characterising the linear and non-linear behavior depicted by elastic Lissajous projections for different R values in oscillatory cycles. (a) depicts $e_1$, the first harmonic, characterising the linear behavior; (b) shows the third harmonic, $e_3$, characterising the non-linear behavior, and (c) shows the third relative harmonic, $e_{3/1}$.

### Section: Cage Modulus

To investigate the softer microstructure due to the formation of bundles, the nature of microstructural changes directly by using a parameter known as the cage modulus ($G_{cage}$) was explored. $G_{cage}$ describes the linear elasticity of the microstructure of the hydrogel network and is evaluated at the point in the rheological response of the hydrogel when there is a momentary mechanical equilibrium within each oscillatory cycle i.e. the elastic and viscous stresses at this point are either zero or momentarily have equal magnitudes with opposite signs [6]. This zero-stress point in the rheological response when examined over increasing $\gamma_0$ is indicative of the changes in the microstructure. The evolution of $G_{cage}$ with respect to R and $\gamma_0$ is presented in Figure S12. At low R values, an extended plateau with normalised $G_{cage}$ of 1 is observeds, aligning with the LVE regime of the hydrogels, suggesting that all the deformation applied is recoverable and no structural changes take place. Interestingly, there is a slight increase in cage modulus at higher $\gamma_0$ just outside the LVE regime which could reflect the irreversible alignment of polymer chains. When $\gamma_0$ is increased beyond the LVE regime, $G_{cage}$ shows a transition to lower values highlighting the structural breakdown of hydrogels leading to softer microstructure. For hydrogels with R < 0.5, the final value of the normalised $G_{cage}$ is approximately 60% of the initial value which suggests that the structural breakdown is partial as the network retains a large part of its integrity. As the R increases beyond 0.5, the nature of the decrease in $G_{cage}$ steadily changes. Specifically, the final value decreases to < 40%, indicating that an increase in R leads to a more catastrophic structural breakdown. At highest R values ( >= 0.67), the evolution of $G_{cage}$ is distinct. The structural rearrangements are observed even at low values of $\gamma_0$ with no distinct plateau in $G_{cage}$ initially. This suggests that hydrogels with high R values are highly capable of undergoing slight microstructural rearrangements even at low $\gamma_0$.

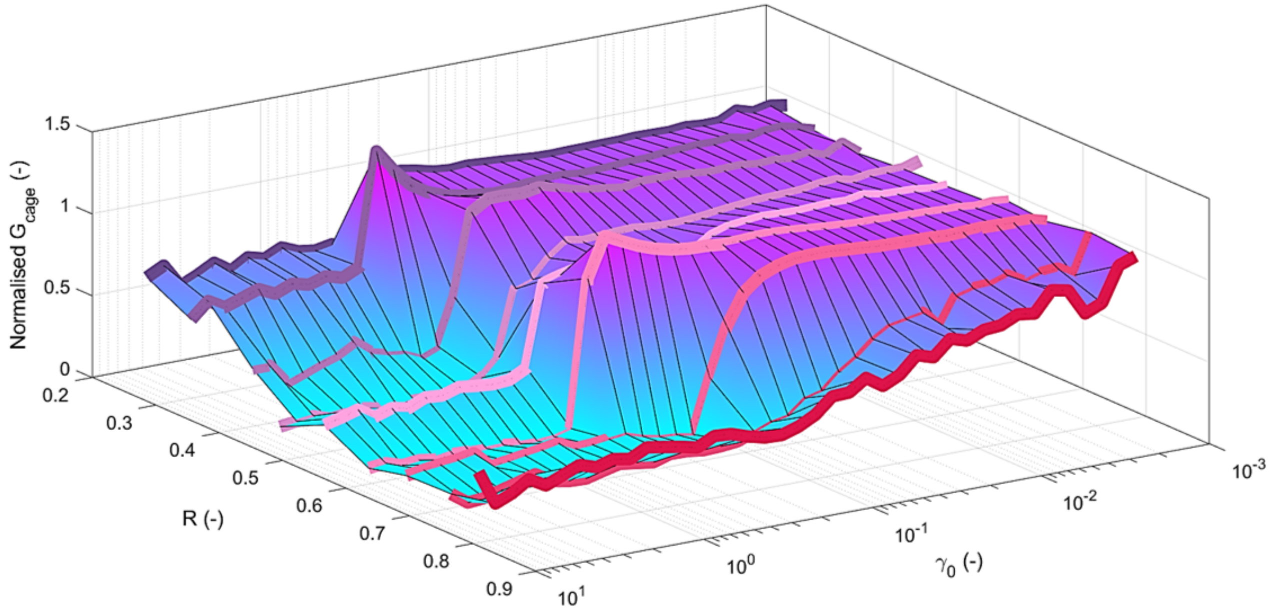


Figure S12: The evolution of normalised cage modulus $G_{cage}$ with increasing $\gamma_0$ for hydrogels with different $Ca^{2+}$ concentration. The cage modulus has been normalised by the average cage modulus observed in the LVE regime of the hydrogels.

**Section: M:G ratio characterization using NMR**

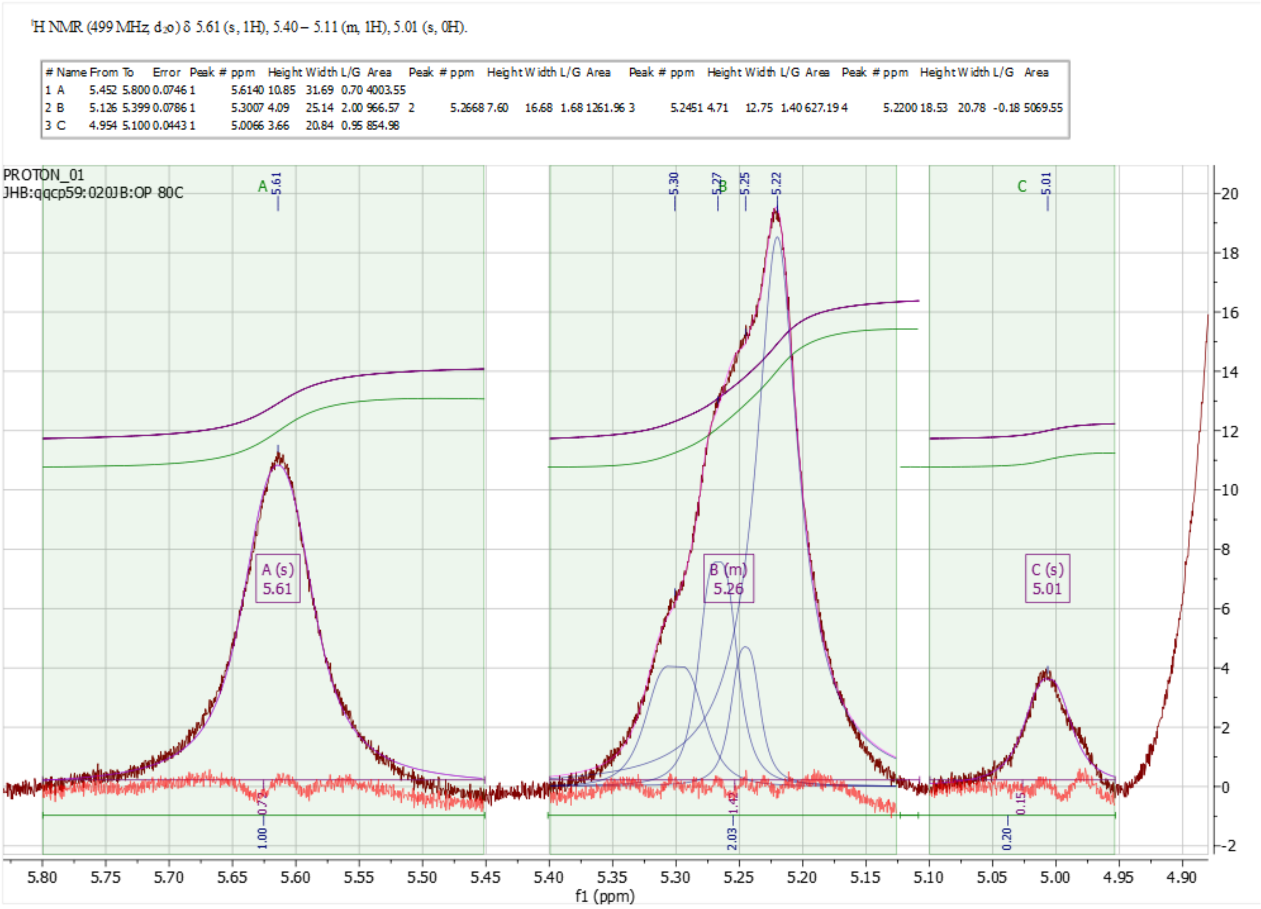


Figure S13: NMR spectra for Dupont Protanal CR8133 sodium alginate

The anomeric proton region of alginate residues (approximately 4.9–5.8 ppm) was analysed to determine the M and G composition. Three main integration regions were identified according to the method proposed by Grasdalen [7] (Table 1).

| | Range | Normalized | Absolute |
|---|---|---|---|
| 1 | 5.80 .. 5.45 | 1 | 3554.96 |
| 2 | 5.40 .. 5.11 | 2.03 | 7217.09 |
| 3 | 5.12 .. 4.95 | 0.2 | 719.75 |

| | Normalized | Absolute |
|---|---|---|
| M/G ratio= | 1.23 | 1.23261 |

Region A corresponds mainly to the H-1 signal of mannuronic acid residues, while regions B and C contain overlapping contributions from guluronic acid and mannuronic acid residues. Due to the partial overlap of signals, the B region was further analysed by peak deconvolution.

The M:G ratio determined by $^1$H NMR was calculated as 1.23.

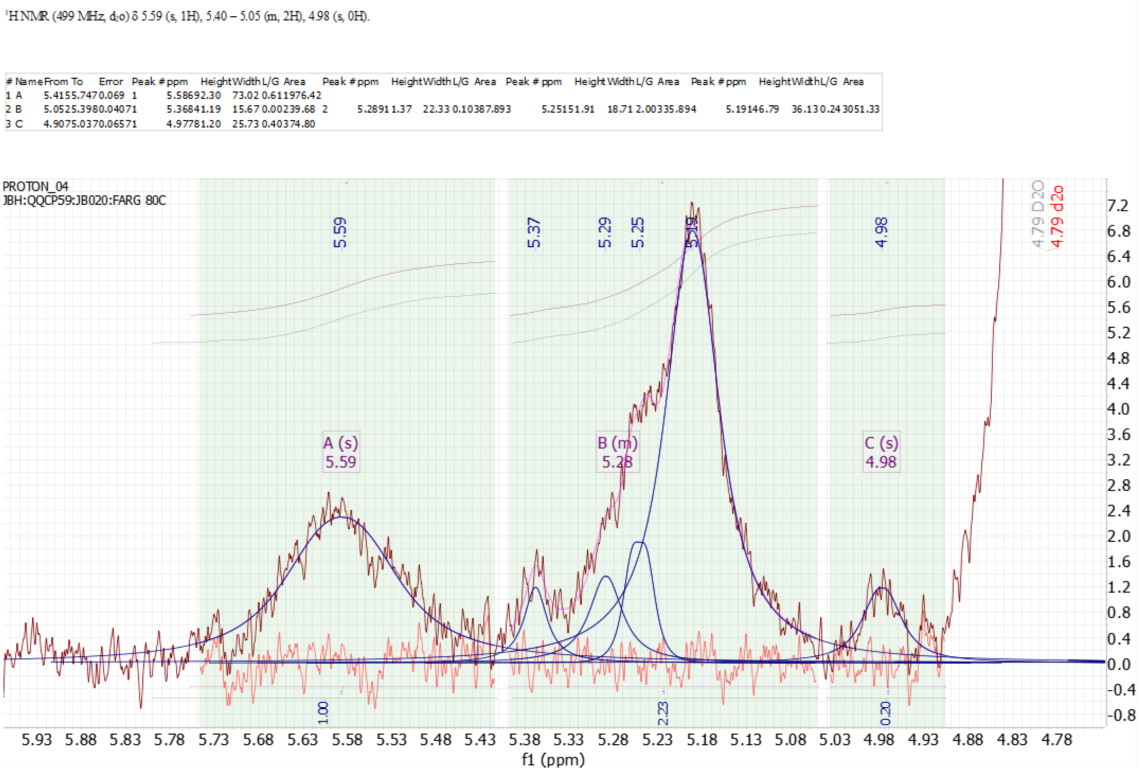


Figure S14: NMR spectra for AL510 sodium alginate

| | Range | Normalized | Absolute |
|---|---|---|---|
| 1 | 5.80 .. 5.41 | 1 | 1591.99 |
| 2 | 5.39 .. 5.05 | 2.23 | 3548.65 |
| 3 | 5.04 .. 4.91 | 0.2 | 312.82 |

| | Normalized | Absolute |
|---|---|---|
| M/G ratio= | 1.43 | 1.42556172 |

For the labelled alginate (Rhodamine-alginate), higher noise is observed in the $^{1}H$ NMR spectrum, which may be associated with the attached fluorophore that can introduce additional structural heterogeneity and reduce the overall spectral quality due to its aromatic groups, potential interactions with the polymer chains, and lower effective concentration of alginate protons contributing to the signal. Furthermore, the conjugation of rhodamine may affect the local chemical environment of nearby protons, resulting in broader peaks and a lower signal-to-noise ratio compared with unlabelled alginate samples. Nevertheless, the characteristic alginate signals (A, B and C) were resolved using the same methodology described for DuPont Protanal CR8133. The evaluated M:G ratio in this case is 1.43.
